# *Spinosaurus* is not an aquatic dinosaur

**DOI:** 10.1101/2022.05.25.493395

**Authors:** Paul C. Sereno, Nathan Myhrvold, Donald M. Henderson, Frank E. Fish, Daniel Vidal, Stephanie L. Baumgart, Tyler M. Keillor, Kiersten K. Formoso, Lauren L. Conroy

## Abstract

A predominantly fish-eating diet was envisioned for the sail-backed theropod dinosaur, *Spinosaurus aegyptiacus*, when its elongate jaws with subconical teeth were unearthed a century ago in Egypt. Recent discovery of the high-spined tail of that skeleton, however, led to a bolder conjecture, that *S. aegyptiacus* was the first fully aquatic dinosaur. The ‘aquatic hypothesis’ posits that *S. aegyptiacus* was a slow quadruped on land but a capable pursuit predator in coastal waters, powered by an expanded tail. We test these functional claims with skeletal and flesh models of *S. aegyptiacus*. We assembled a CT-based skeletal reconstruction based on the fossils, to which we added internal air and muscle to create a posable flesh model. That model shows that on land *S. aegyptiacus* was bipedal and in deep water was an unstable, slow surface swimmer (<1m/s) too buoyant to dive. Living reptiles with similar spine-supported sails over trunk and tail in living reptiles are used for display rather than aquatic propulsion, and nearly all extant secondary swimmers have reduced limbs and fleshy tail flukes. New fossils also show that *Spinosaurus* ranged far inland. Two stages are clarified in the evolution of *Spinosaurus*, which is best understood as a semiaquatic bipedal ambush piscivore that frequented the margins of coastal and inland waterways.

## Introduction

In 1915 Ernst von Stromer announced the discovery in Egypt’s Western Desert of the elongate jaws and partial skeleton of a large sail-backed predator, *Spinosaurus aegyptiacus* (***Stromer, 1915***). Other bones found nearby (***Stromer, 1934***) contributed to Stromer’s (***1936***) initial reconstruction of *S. aegyptiacus* as a sail-backed, piscivorous biped, shortly before all of these bones were destroyed in WWII (***Nothdurft and Smith, 2002***; ***Smith et al., 2006***). Over the last 30 years, additional skull and postcranial bones came to light in western Morocco in beds of similar age to those in Egypt (***Russell, 1996***; ***Dal Sasso et al., 2005***; ***Smyth et al., 2020***; ***Ibrahim et al., 2020b***). Central among these finds was a partial skeleton (designated the neotype) that allowed a more complete reconstruction, confirming its interpretation as a semiaquatic piscivore (***Ibrahim et al., 2014***).

As skeletal information on the unusual predator improved, so has speculation as to whether *S. aegyptiacus* was better adapted to life in water as an aquatic predator, based on inferences from oxygen isotopes in enamel (***Amiot et al., 2010***), the dental rosette likened to the jaws of a conger eel (***Vullo et al., 2016***), the alleged elevated positioning of the orbits in the skull for visibility while largely submerged (***Arden et al., 2019***), the hypothetical underwater role of the trunk sail (***Gimsa et al., 2016***), and the infilling of the medullary cavities of hind limb bones that may have functioned as ballast (***Ibrahim et al., 2014***; ***Aureliano et al., 2018***).

### The aquatic hypothesis

Recent discovery of the tall-spined tail bones of the neotypic skeleton reinvigorated the interpretation of *S. aegyptiacus* as the first fully aquatic dinosaur (***Ibrahim et al., 2020a***) here dubbed the ’aquatic hypothesis,’ which makes three basic propositions. Unlike any other theropod, according to the hypothesis, *S. aegyptiacus* (1) reverted to a quadrupedal stance on land, ostensibly knuckle-walking with long-fingered, long-clawed forelimbs, while (2) functioning in water as a capable, diving pursuit predator, using an expanded tail as a “novel propulsor organ”. Its adaptations for “deep-water propulsion” were reasserted by quantitative analysis of bone density that suggested *S. aegyptiacus* was a frequent diver (***Fabbri et al., 2022***), an interpretation that has been challenged (***Myhrvold et al., in press***). A deep-diving pursuit predator of large body size, furthermore, would be found exclusively in (3) deep-water coastal or marine habitats rather than also ranging into freshwater, inland environments. We test these three central propositions.

Critique of the aquatic hypothesis thus far has focused on an alternative functional explanation for the high-spined tail (as a display structure) and qualitative functional interpretations of its skeletal anatomy (***Hone and Holtz, 2021***). Biomechanical evaluation of the aquatic functionality of *S. aegyptiacus* remains rudimentary. The propulsive capacity of the tail in water was judged to be better than terrestrial counterparts by oscillating miniature plastic tail cut-outs in water (***Ibrahim et al., 2020a***), a limited approximation of the biomechanical properties of an anguilliform tail (***Lighthill, 1969***; ***van Rees et al., 2013***; ***Gutarra and Rahman, 2022***) and failed to take into account the bizarre anterior half of the animal. The center of body mass, a critical functional parameter, has been estimated for *S. aegyptiacus* three times, each estimate pointing to a different location ranging from the middle of the trunk to a position over the hind limbs (***Henderson, 2018***; ***Ibrahim et al., 2014, 2020a***). Quantitative comparisons have not been made regarding the size or surface area of the limbs, hind feet and tail of *S. aegyptiacus* to counterparts in extant primary or secondary swimmers.

Adequate evaluation of the aquatic hypothesis, thus, requires more realistic biomechanical tests, quantitative body, axial and limb comparisons between *S. aegyptiacus* and extant primary and secondary swimmers, and a survey of bone structure beyond the femur and shaft of a dorsal rib. Such tests and comparisons require an accurate 3D digital flesh model of *S. aegyptiacus*, which, in turn, requires an accurate skeletal model. Hence, we began this study by assembling a complete set of CT scans of the fossil bones for *S. aegyptiacus* and its African forerunner, *Suchomimus tenerensis* (***Sereno et al., 1998***).

### Aquaphilic terminology

Aquatic status is central to the ‘aquatic hypothesis.’ *S. aegyptiacus*, the hypothesis holds, is the first non-avian dinosaur bearing skeletal adaptations devoted to lifestyle and locomotion in water, some of which inhibited terrestrial function. The contention is that *S. aegyptiacus* was not only a diving pursuit predator in the open water column, but also a quadruped on land with long-clawed forelimbs poorly adapted for weight support. A later publication seemed to downgrade that central claim by suggesting that any vertebrate with “aquatic habits,” such as wading, submergence or diving, had an “aquatic lifestyle” (***Fabbri et al., 2022***). That broadened usage of “aquatic lifestyle,” however, blurs the longstanding use of aquatic as applied to lifestyle (***Pacini and Harper, 2008***). We outline below traditional usage of aquaphilic terms, which we follow.

The adjective “aquatic” is used either as a broad categorization of *lifestyle* or, in more limited capacity, in reference to an *adaptation* of a species or group. In the former case, a vertebrate with an “aquatic lifestyle” or “aquatic ecology” is adapted for life primarily, or solely, in water with severely reduced functional capacity on land (***Pacini and Harper, 2008***). *Aquatic vertebrates* (e.g., bony fish, sea turtles, whales) live exclusively or primarily in water and exhibit profound cranial, axial or appendicular modifications for life in water, especially at larger body sizes (***Webb, 1984***; ***Webb and De Buffrénil, 1990***; ***Hood, 2020***). For example, extant whales are secondarily aquatic mammals that spend all of their lives at sea and exhibit profound skeletal modifications for aquatic sensory and locomotor function. A marine turtle, similarly, is considered an aquatic reptile, regardless of whether it clambers ashore briefly to lay eggs, because the vast majority of its life is spent in water using profoundly modified limbs for aquatic locomotion (flippers) that function poorly on land.

An aquaphilic animal with less profound adaptations to an aqueous arena is said to be *semi-aquatic* (or semi-aquatic), no matter the proportion of aquatic foodstuffs in its diet, the proportion of time spent in water, or the proficiency of swimming or diving. Nearly all semiaquatic vertebrates are secondarily aquaphilic, having acquired aquatic adaptations over time to enhance functional capacity in water without seriously compromising terrestrial function (***Howell, 1930***; ***Hood, 2020***). Indeed, semiaquatic animals are also semiterrestrial (***Fish, 2016***). Freshwater turtles, for example, are regarded as semiaquatic reptiles because they frequent water rather than live exclusively within an aqueous habitat, are sometimes found in inland habitats, and exhibit an array of less pro-found modifications (e.g., interdigital webbing) for locomotion in water (***Pacini and Harper, 2008***). Extant crocodylians and many waterbirds, likewise, are capable swimmers and divers but retain excellent functional capacity on land. Auks (Alcidae), among the most water-adapted of semiaquatic avians, are agile wing-propelled, pursuit divers with an awkward upright posture on land resembling penguins, but they retain the ability to fly and inhabit land for extended periods (***Nettleship, 1996***). On the other hand, the flightless penguins (Sphenisciformes) are considered aquatic due to their more profound skeletal modifications for swimming and deep diving and more limited terrestrial functionality, although still retaining the capability to trek inland and stand for considerable durations while brooding. As nearly all semiaquatic vertebrates have an aquatic diet and the ability to swim or dive, more profound functional allegiance to water is requisite for an “aquatic” appellation (***Pacini and Harper, 2008***).

An *aquatic adaptation* of an organism refers to the function of a particular feature, not the overall lifestyle of an organism. That feature should have current utility and primary function in water (***Houssaye and Fish, 2016***). Aquatic adaptations are presumed to have evolved their functionality in response to water and cannot also have special functional utility in a subaerial setting. For example, the downsized, retracted external nares in *S. aegyptiacus* would prevent water intake through the nostrils while feeding with the snout submerged. There has yet been offered no plausible alternative explanation involving terrestrial function for the downsizing and retraction of the external nares in spinosaurids, a unique condition among non-avian theropods. The hypertrophied neural spines of the tail of *S. aegyptiacus*, in contrast, are ambiguous as an “aquatic adaptation,” because expanded tails can function both as aquatic propulsors and terrestrial display structures. For the expanded tail to be an “aquatic adaptation,” its morphological construction and biomechanical function demonstrate its primary utility and capability in water, as shown in extant tail-powered primarily or secondarily aquatic vertebrates (e.g., newts, crocodylians, beavers, otters; (***Fish et al., 2021***). The same must be shown or inferred to be the case in extinct secondarily aquatic vertebrates (***Gutarra and Rahman, 2022***). We have not found such substantiating evidence in tail form and inferred function in *S. aegyptiacus* or other spinosaurids for the heightened tail to be substantiated as an aquatic adaptation (see below).

### Our approach

To test the aquatic hypothesis for *S. aegyptiacus*, we began with CT scans of spinosaurid fossils from sites in Africa to build high-resolution 3D skeletal models of *S. aegyptiacus* (***Figure 1****A*) and its fore-runner *Suchomimus tenerensis* (***Figure 1****F*). Many vertebrae and long bones in both genera show significant internal pneumatic (air) or medullary (marrow) space, which has ramifications for buoyancy. When compared to the 2D silhouette drawing used in the aquatic hypothesis (***Ibrahim et al., 2020a***), our CT-based 3D skeletal model of *S. aegyptiacus* differs significantly in skeletal proportions.

**Figure 1.**
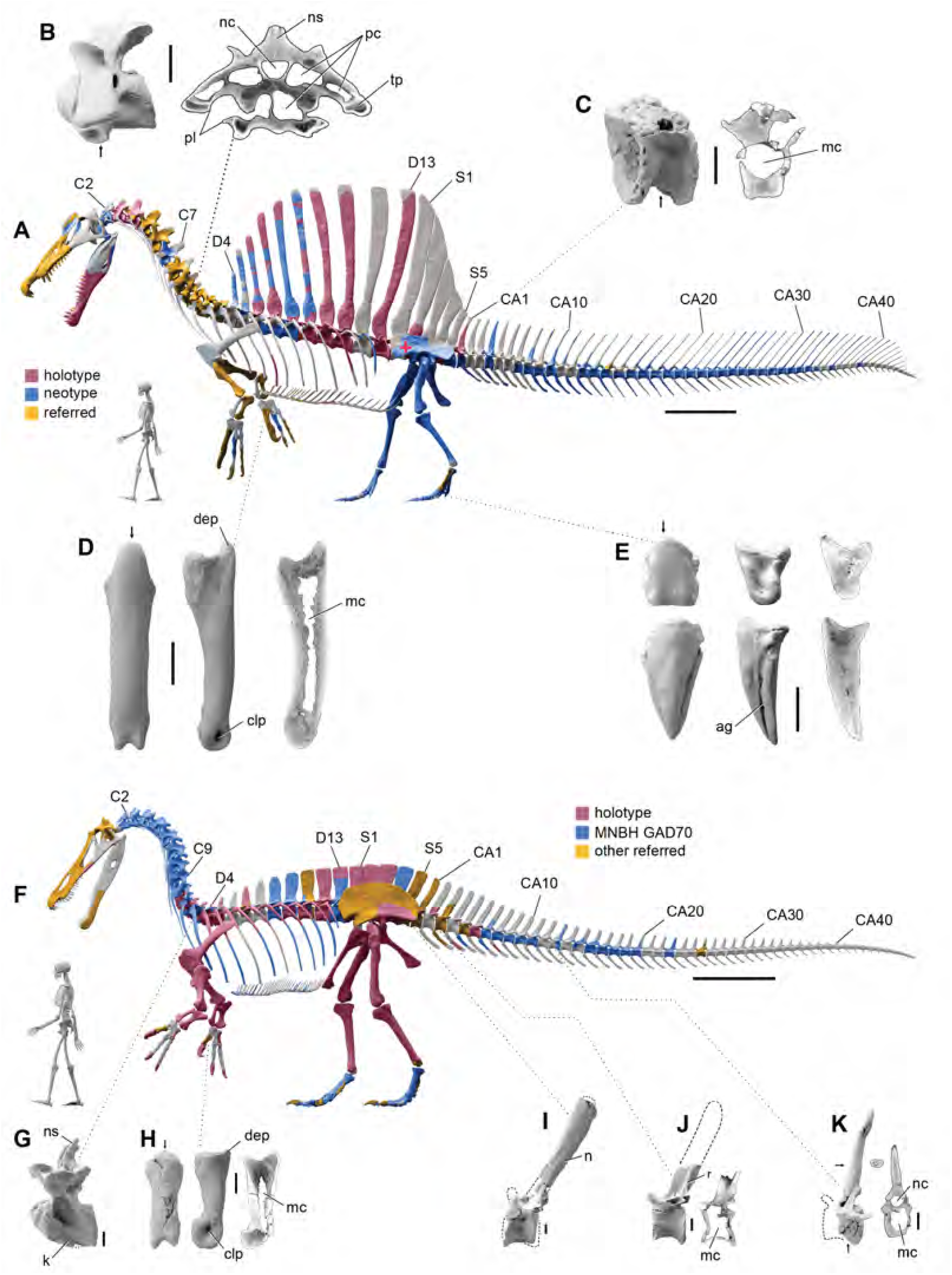
Digital skeletal reconstructions of the African spinosaurids *Spinosaurus aegyptiacus* and *Suchomimus tenrensis*. **(A)** *Spinosaurus aegyptiacus* (early Late Cretaceous, Cenomanian, ca. 95 Ma) showing known bones based on the holotype (BSPG 1912 VIII 19, red), neotype (FSAC-KK 11888, blue), and referred specimens (yellow) and the center of mass (cross) of the flesh model in bipedal stance (overlap priority: neotype, holotype, referred bones). **(B)** Cervical 9 (BSPG 2011 I 115) in lateral view and coronal cross-section showing internal air space. **(C)** Caudal 1 centrum (FSAC-KK 11888) in anterolateral view and coronal CT cross-section. **(D)** Right manual phalanx I-1 (UCRC PV8) in dorsal, lateral and sagittal CT cross-sectional views. **(E)** Pedal phalanges IV-4, IV-ungual (FSAC-KK 11888) in dorsal, lateral and sagittal CT. **(F)** *Suchomimus tenerensis* (mid Cretaceous, Aptian-Albian, ca. 110 Ma) showing known bones based on the holotype (MNBH GAD500, red), a partial skeleton (MNBH GAD70, blue), and other referred specimens (yellow) and the center of mass (cross) of the flesh model in a bipedal stance (overlap priority: holotype, MNBH GAD70, referred bones). **(G)** Dorsal 3 in lateral view (MNBH GAD70). **(H)** Left manual phalanx I-1 (MNBH GAD503) in dorsal, lateral and sagittal CT cross-sectional views. **(I)** Caudal 1 vertebra in lateral view (MNBH GAD71). **(J)** Caudal ∼3 vertebra in lateral view (MNBH GAD85). **(K)** Caudal ∼13 vertebra in lateral view with CT cross-sections (coronal, horizontal) of the hollow centrum and neural spine (MNBH GAD70). ag, attachment groove; C2, 7, 9, cervical vertebra 2, 7, 9; CA1, 10, 20, 30, 40, caudal vertebra 1, 10, 20, 30, 40; clp, collateral ligament pit; D4, 13, dorsal vertebra 4, 13; dip, dorsal intercondylar process; k, keel; mc, medullary cavity; nc, neural canal; ns, neural spine; pc, pneumatic cavity; pl, pleurocoel; r, ridge; S1, 5, sacral vertebra 1, 5. Dashed lines indicate contour of missing bone, arrows indicate plane of CT-sectional views, and scale bars equal 2 m (A, F), 5 cm (B, C), 3 cm (D, E, H-K) with human skeletons 1.8 m tall (A, F).

We enveloped the skeletal model in flesh informed by CT scans revealing the muscle volume and air spaces in extant reptilian and avian analogs. To create a 3D flesh model for *S. aegyptiacus* (***Figure 2****A, B*), internal air spaces (trachea, lungs, air sacs) were shaped and positioned as in extant analogs. We created three options for internal air volume based on extant squamate, crocodilian and avian conditions (***Figure 2****C*) and assigned densities to body partitions based on local tissue types and air space. We calculated the surface area and volume of the flesh model as well as its component body parts.

**Figure 2.**
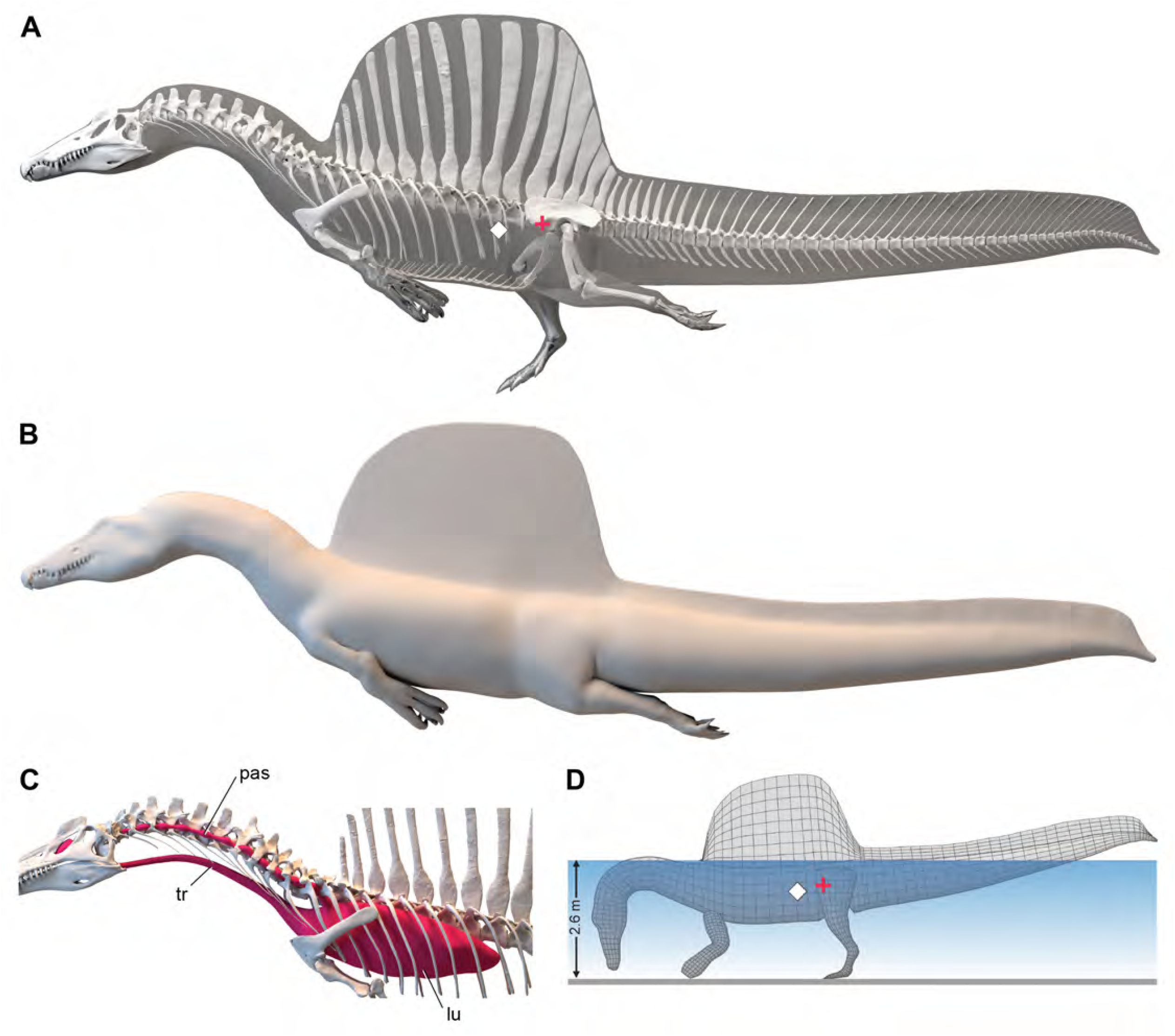
Digital flesh model of *Spinosaurus aegyptiacus*. **(A)** Translucent flesh model in hybrid swimming pose showing centers of mass (cross) and buoyancy (diamond). **(B)** Opaque flesh model in axial swimming pose with adducted limbs. **(C)** Modeled air spaces (“medium” option) include pharynx-trachea, lungs and paraxial air sacs. **(D)** Wading-strike pose at the point of flotation (2.6 m water depth) showing centers of mass (cross) and buoyancy (diamond). Abbreviations: lu, lungs; pas, paraxial air sacs; tr, trachea.

We posed this integrated flesh model in bipedal, hybrid- and axial-powered poses, the latter two based on the swimming postures of extant semiaquatic reptiles (***Grigg and Kirshner, 2015***, Figure 2B). We calculated *center of mass* (CM) and *center of buoyancy* (CB) to evaluate the habitual two- or four-legged stance of *S. aegyptiacus* on land (***Figure 1****A*), the depth of water at the point of flotation (***Figure 2****D*), and the neutral position of the flesh model in deeper water (***Figure 2****A, B*). Using biomechanical formulae (***Lighthill, 1969***) and data from extant alligators (***Fish, 1984***), we estimated the maximum force output of its tail, which was used to calculate maximum swimming velocity at the surface and underwater (***Figure 3****A*). We also evaluated its stability, maneuverability and diving potential in water (***Figure 3****B*), with all of these functional capacities compared to extant large-bodied aquatic vertebrates.

**Figure 3.**
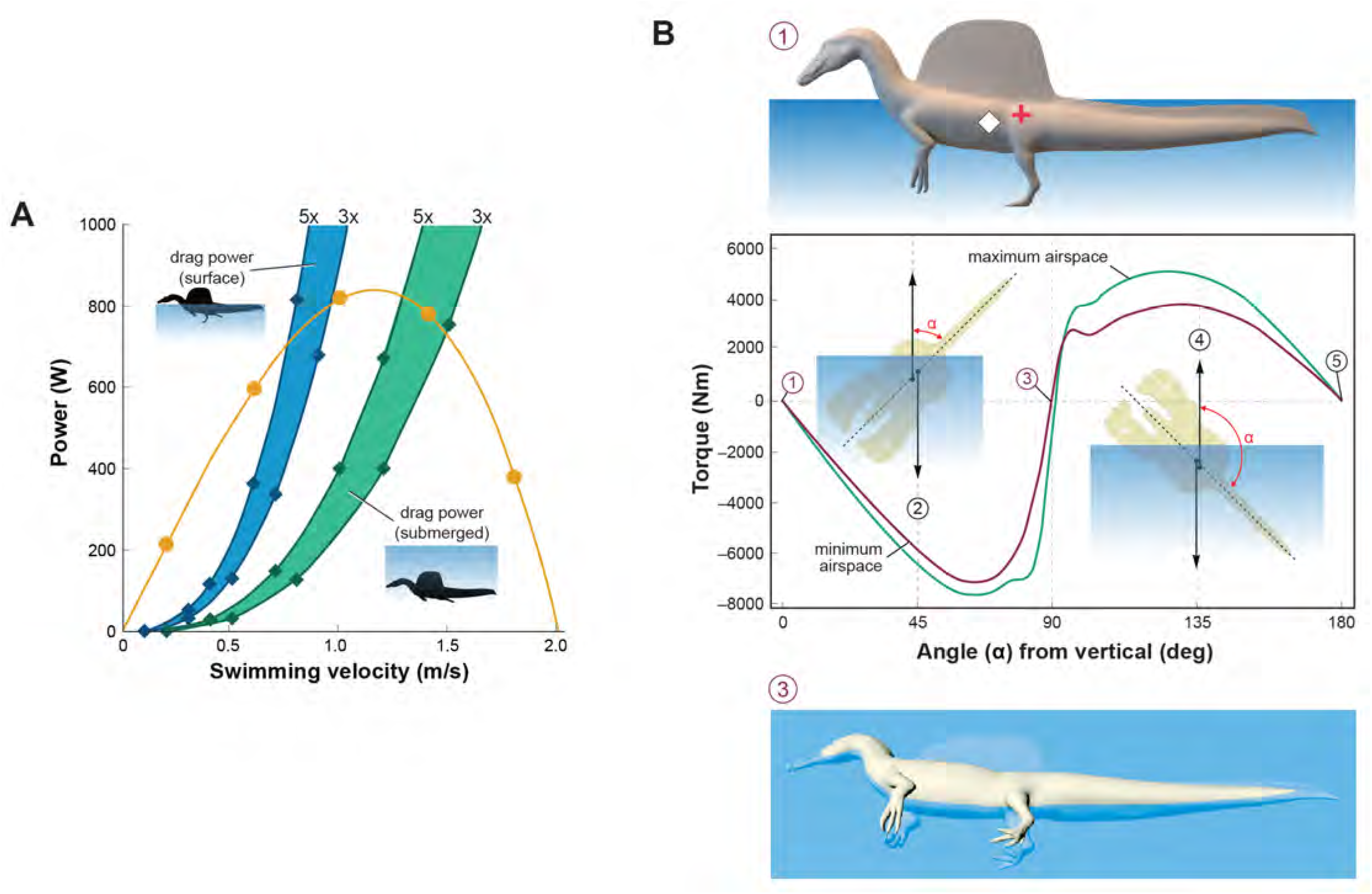
Biomechanical evaluation of *S. aegyptiacus* in water. **(A)** Tail thrust (yellow curve) and opposing drag forces as a function of swimming velocity at the surface (blue) and submerged (green), with drag during undulation estimated at 3 and 5 times stationary drag. **(B)** Stability curve for the flesh model of *S. aegyptiacus* in water showing torque between the centers of mass (cross) and buoyancy (diamond), unstable equilibria when upright or upside down (positions 1, 5), and a stable equilibrium on its side (position 3) irrespective of the volume of internal air space. Curves are shown for flesh models with minimum (magenta) and maximum (green) air spaces with a dashed line showing the vertical body axis and vector arrows for buoyancy (up) and center of mass (down).

We turned to extant analogs to consider the structure and function of similar spine-supported sails over the trunk and tail in lizards and the form of tail vertebrae in tail-powered secondary swimmers (***Figure 4***). We also considered the relative size (surface area) of appendages in a range of secondary swimmers (***Figure 5***), and how the surface area of foot paddles and tail scale in crocodylians (***Figure 6***).

**Figure 4.**
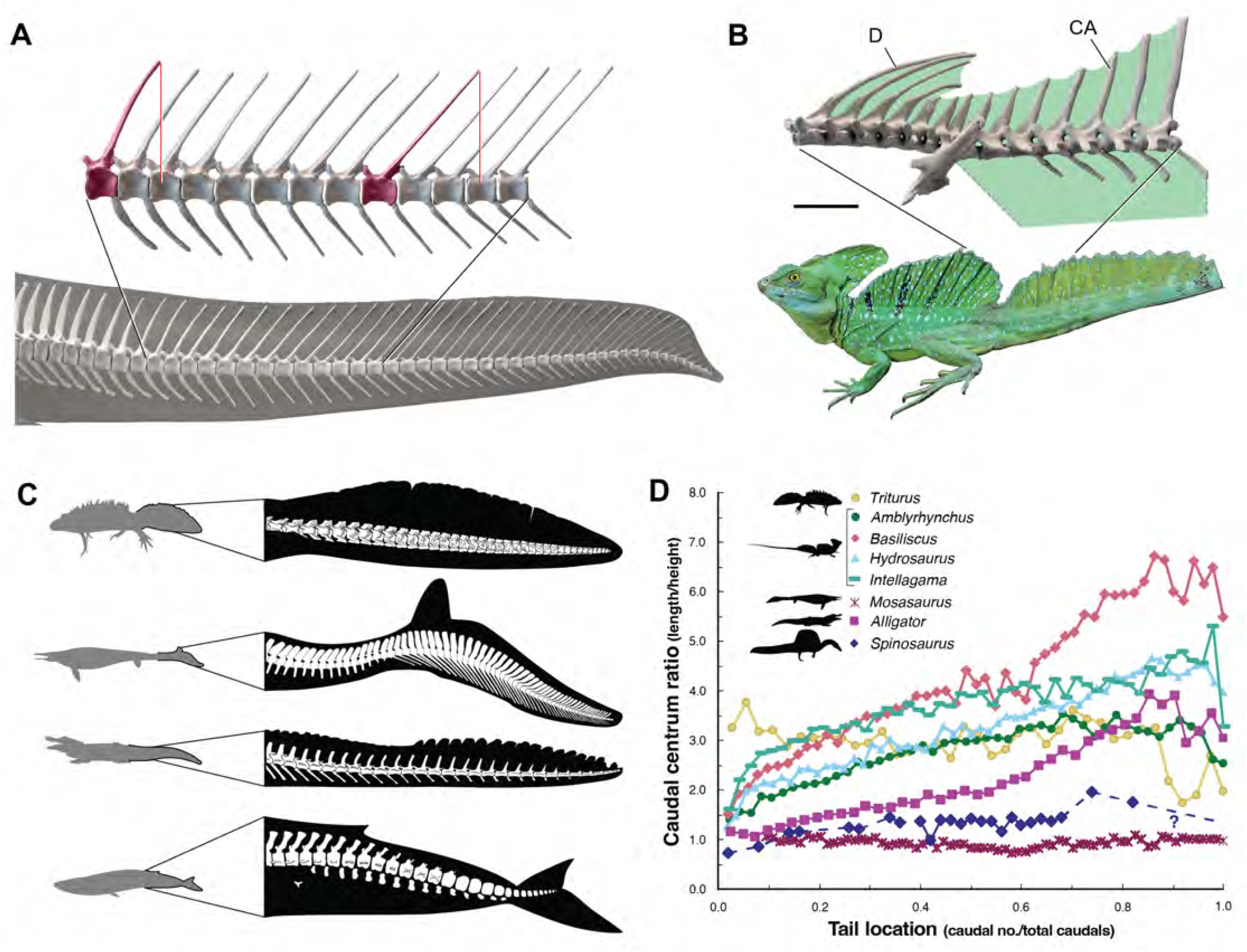
Skeletal comparisons between *S. aegyptiacus*, a basilisk lizard and secondarily aquatic vertebrates. **(A)** Tail in *S. aegyptiacus* showing overlap of individual neural spines (red) with more posterior vertebral segments. **(B)** Sail structure in the green basilisk (CT-scan enlargement) and in vivo form and coloration of the median head crest and sail (*Basiliscus plumifrons* FMNH 112993). **(C)** Tail flukes in a newt, mosasaur, crocodilian and whale. **(D)** Centrum proportions along the tail in the northern crested newt (*Triturus cristatus* FMNH 48926), semiaquatic lizards (marine iguana *Amblyrhynchus cristatus* UF 41558, common basilisk *Basiliscus basiliscus* UMMZ 121461, Australian water dragon *Intellagama lesueurii* FMNH 57512, sailfin lizard *Hydrosaurus amboinensis* KU 314941), an extinct mosasaurid (*Mosasaurus* sp. UCMP 61221; ***Lindgren et al. (2013***)), an alligator (*Alligator mississippiensis* UF 21461), and *Spinosaurus* (*S. aegyptiacus* FSAC-KK 11888). Data in ***Appendix 2***.

**Figure 5.**
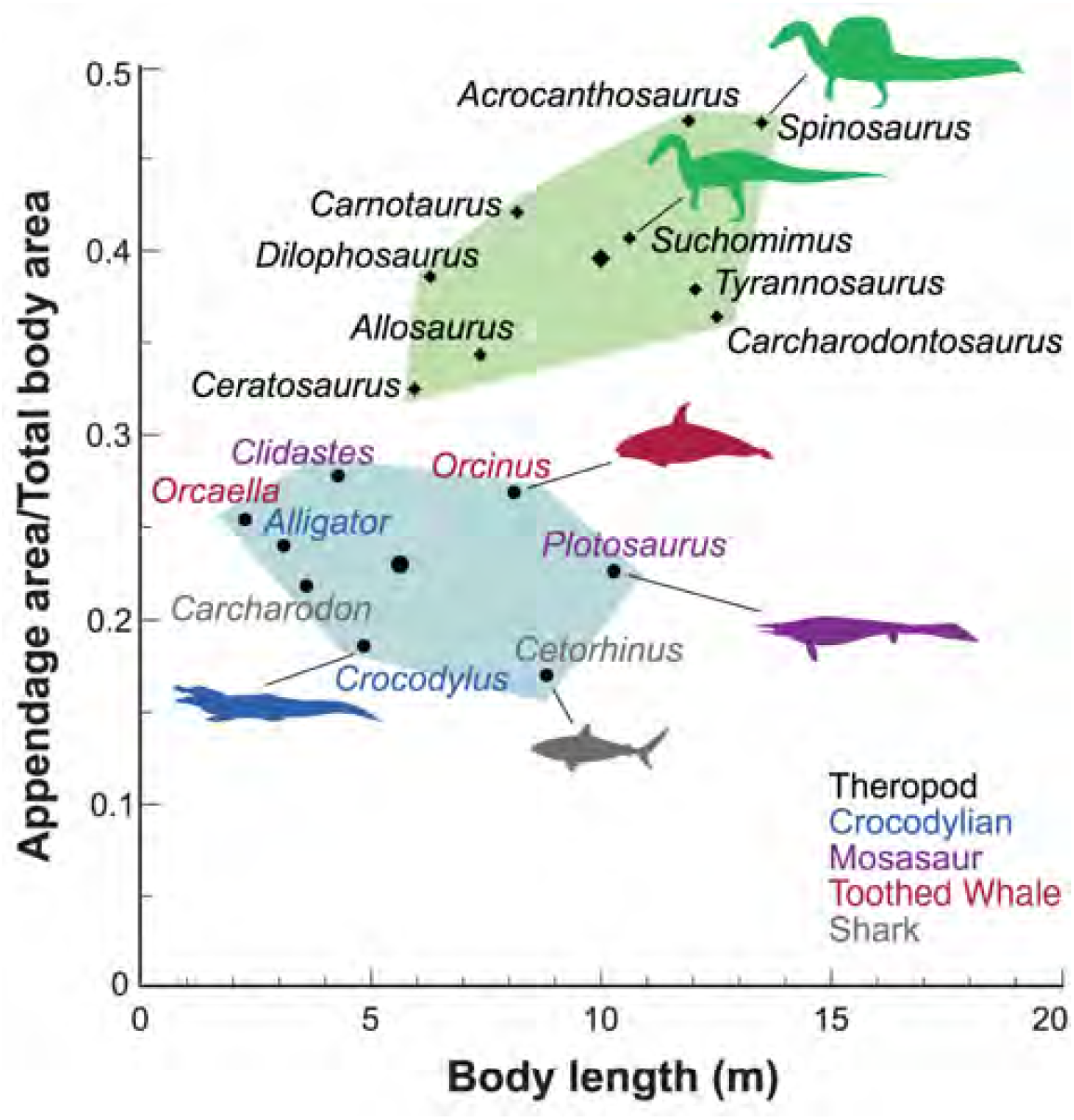
Appendage versus total body surface area in aquatic and semiaquatic vertebrates. *Spinosaurus aegyptiacus* and other non-avian theropods (green hull, diamonds) have appendages with considerable surface area compared to aquatic and semiaquatic vertebrates (blue hull, circles).

**Figure 6.**
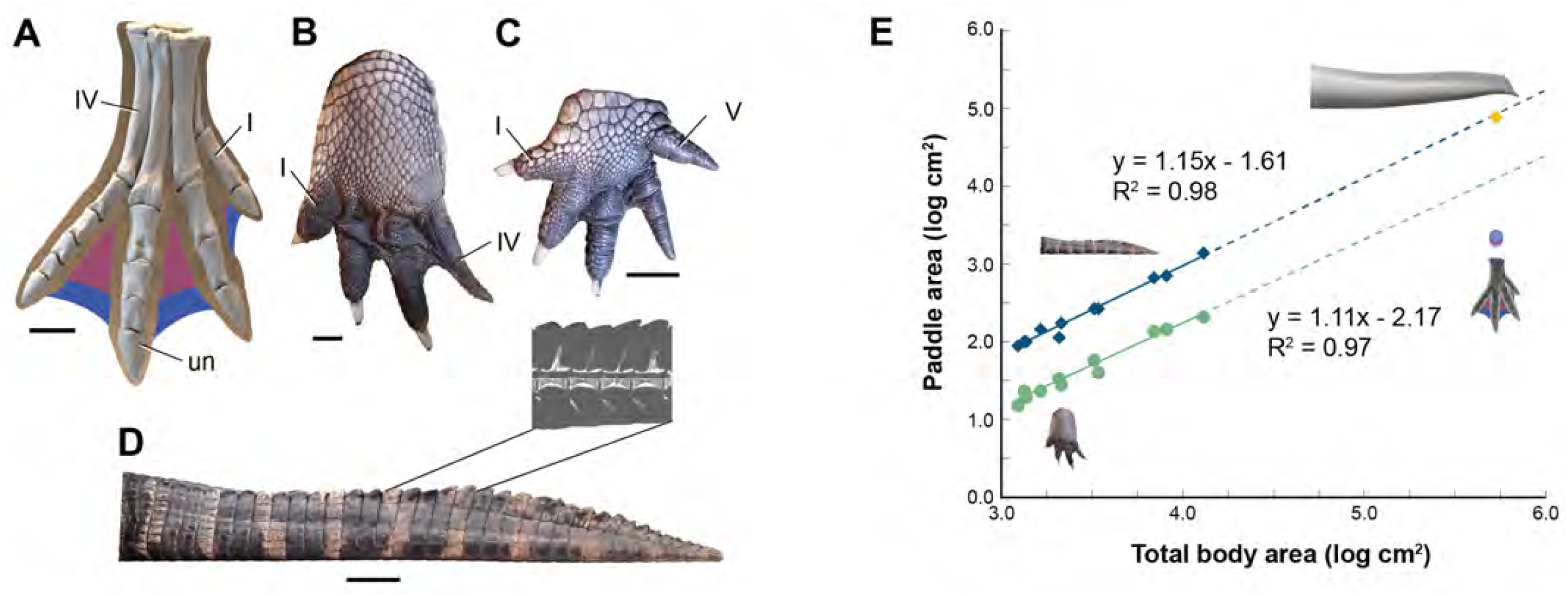
Appendage surface area and scaling of paddle surface areas in crocodylians. **(A)** Right hind foot of *S. aegyptiacus* (FSAC-KK 11888) showing the outlines of digital flesh based on the living ostrich (*Struthio camelus*) as well as partial (pink) and full (blue) interdigital webbing. **(B)** Hindfoot of an adult *Alligator mississippiensis* (WDC) in ventral view. **(C)** Forefoot of an adult *Alligator mississippiensis* (WDC) in ventral view. **(D)** Tail of an adult *Alligator mississippiensis* (WDC) in lateral view with CT visualization of vertebrae within the fleshy tail fluke. **(E)** Log-log plot of surface areas of webbed hindfoot and side of the tail as a function of total body area in a growth series for *Alligator mississippiensis* (hindfoot, green dots; tail, blue diamonds) and adult *S. aegyptiacus* (hindfoot, purple-blue dots; tail, yellow diamond). Abbreviations: I, IV, V digits I, IV, V, un, ungual. Scale bars are 10 cm (A) and 3 cm (B-D).

Lastly, we turned to the spinosaurid fossil record to look at the habitats where spinosaurid fossils have been found. We reviewed their distribution (***Figure 7***) to determine if spinosaurids, and *S. aegyptiacus* in particular, were restricted to coastal, marine habitats like all large secondarily aquatic vertebrates. We updated spinosaurid phylogeny in order to discern major stages in the evolution of spinosaurid piscivorous adaptations and sail structures (***Figure 8***), incorporating the latest finds including new fossils of *Spinosaurus* from Niger.

**Figure 7.**
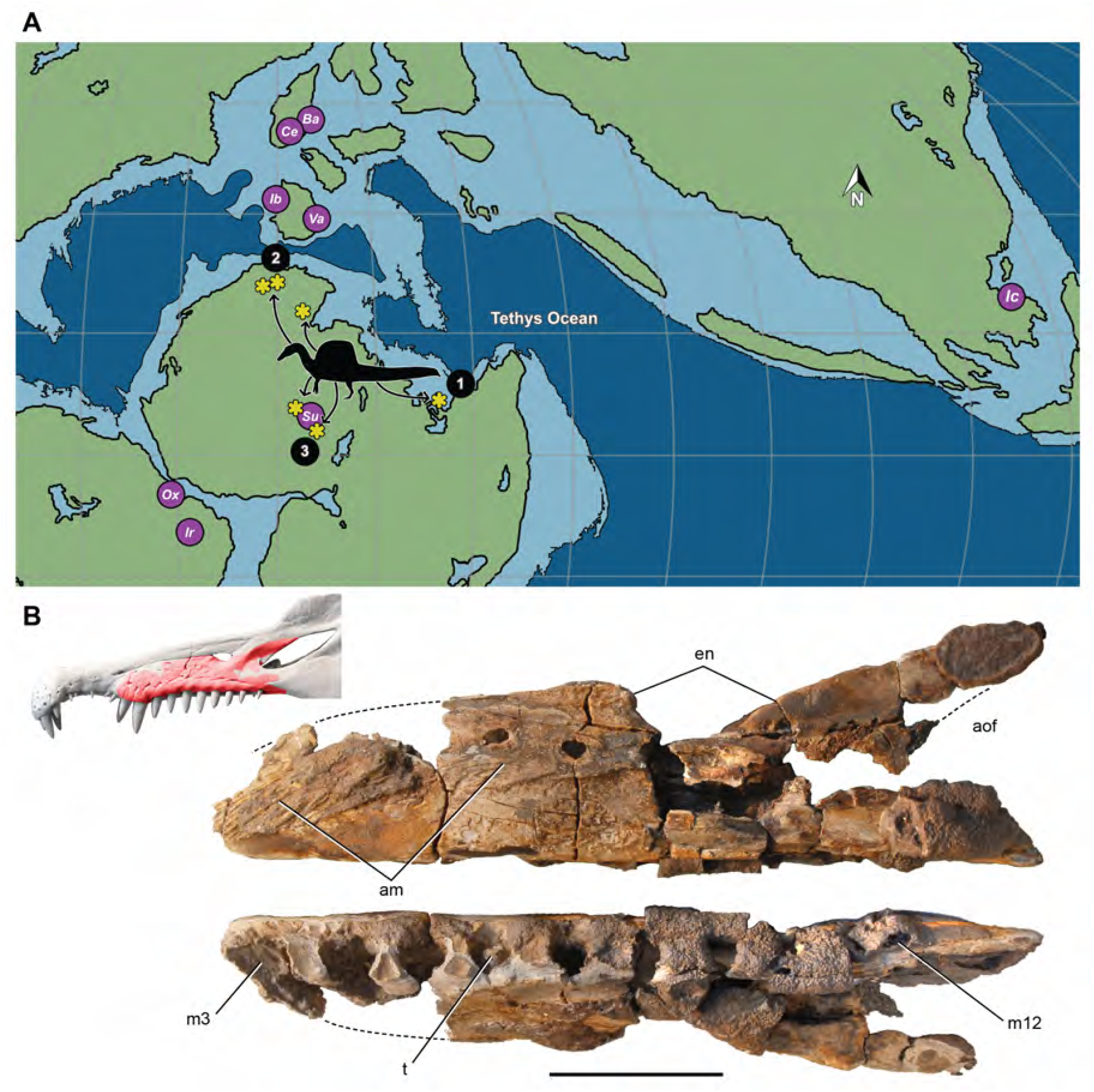
Paleogeographic location of spinosaurid fossils. **(A)** Paleogeographic map (early Albian, ∼110 Mya; ***Scotese (2014***)). showing the circum-Tethyan fossil localities for baryonychines (*Baryonyx, Suchomimus*) and spinosaurines (*Ichthyovenator, Vallibonavenatrix, Oxalaia, Irritator/Angaturama, Spinosaurus*). *Spinosaurus* localities (yellow asterisks) range across northern Africa from coastal (sites 1, 2) to inland (site 3) sites. **(B)** *Spinosaurus* sp. right maxilla (MNBH EGA1) from Égaro North (central Niger) in medial (top) and ventral (bottom) views and shown (red) superposed on the snout of *Spinosaurus aegyptiacus*. Abbreviations: 1, *S. aegyptiacus* holotype (Bahariya, Egypt); 2, *S. aegyptiacus* neotype (Zrigat, Morocco); 3, *Spinosaurus* sp. (Égaro North, Niger); am, articular rugosities for opposing maxilla; aofe, antorbital fenestra; Ba, *Baryonyx walkeri*; en, external naris; Ic, *Ichthyovenator laosensis*; Ir, *Irritator challenger/Angaturama limai*; m3, 12, maxillary tooth alveolus 3, 12; Ox, *Oxalaia quilombensis*; Su, *Suchomimus tenerensis*; t, tooth; Va, *Vallibonavenatrix cani*. Scale bar is 10 cm.

**Figure 8.**
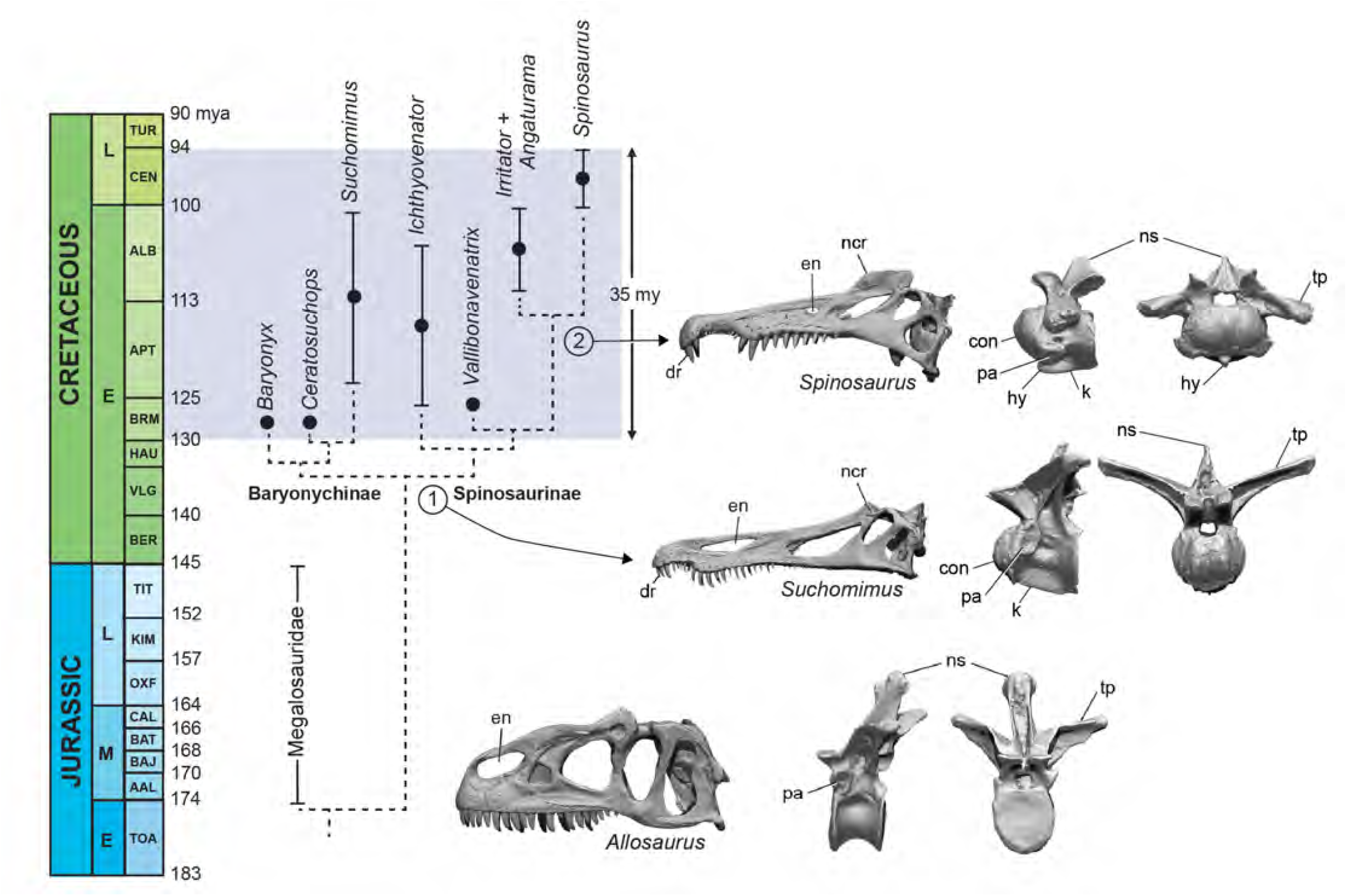
Calibrated phylogeny of spinosaurids (Barremian to Cenomanian, ∼35 My). Updated phylogenetic analysis of spinosaurids resolves two stages in the evolution of piscivory and display. We show key cranial adaptations in the skull and highlight changes at the anterior end of the trunk to enhance neck ventroflexion (second dorsal vertebra in lateral and anterior views). Bottom, the fully terrestrial theropod *Allosaurus fragilis* (***Madsen, 1976***); middle, the baryonychine spinosaurid *Suchomimus tenerensis* (MNBH GAD70); top, the spinosaurine *Spinosaurus aegyptiacus* (BSPG 1912 VIII 19). Abbreviations: con, condyle; dr, dental rosette; en, external naris; hy, hypopophysis; k, keel; ncr, nasal crest; ns, neural spine; pa, parapophysis; tp, transverse process.

### Institutional abbreviations

BSPG, Bayerische Staatssammlung für Paläontologie und Geologie, Munich, Germany; FMNH, Field Museum of Natural History, Chicago, USA; FSAC, Faculté des Sciences Aïn Chock, University of Casablanca, Casablanca, Morocco; KU, The University of Kansas, Museum of Natural History, Lawrence, USA; MNBH, Musée National de Boubou Hama, Niamey, Niger; MNHN, Muséum national d’Histoire naturelle, Paris, France; NMC, Canadian Museum of Nature, Ottawa, Canada; UCMP, University of California, Museum of Paleontology, Berkeley, USA; UCRC, University of Chicago Research Collection, Chicago, USA; UF, University of Florida, University of Florida Collections, Gainesville, USA; UMMZ, University of Michigan, Museum of Zoology, Ann Arbor, USA; WDC, Wildlife Discovery Center, Lake Forest, USA.

## Results

### Spinosaurid skeletal models

Our skeletal reconstruction of *S. aegyptiacus* is just under 14 m long (***Figure 1****A*), which is more than 1 m shorter than previously reported (***Ibrahim et al., 2014***). Major differences are apparent when compared to the 2D graphical reconstruction of the aquatic hypothesis (***Ibrahim et al., 2020a***). In their reconstruction, the length of the presacral column, depth of the ribcage, and length of the forelimb were overestimated by approximately 10%, 25% and 30%, respectively, over dimensions based on CT-scanned fossils. When translated to a flesh model, all of these proportional over-estimates (heavier neck, trunk, forelimb) shift the center of mass anteriorly (see Materials and Methods).

The hind limb long bones (femur, tibia, fibula, metatarsals) in *S. aegyptiacus* lack the medullary cavity common to most dinosaurs and theropods in particular. When first discovered, the infilled hind limb bones in *S. aegyptiacus* were interpreted as ballast for swimming (***Ibrahim et al., 2014***). The infilled condition, however, is variable as shown by the narrow medullary cavity in a femur of another individual slightly larger than the neotype (***Myhrvold et al., in press***). The bone infilling, furthermore, is fibrolamellar and cancellous, similar to the infilled medullary cavities of other large-bodied terrestrial dinosaurs (***Vanderven et al., 2014***) and mammals (***Houssaye et al., 2016***). In contrast, dense pachystotic bone composes the solid and sometimes swollen bones of some secondarily aquatic vertebrates that use increased skeletal mass as ballast (***Houssaye, 2009***).

Medullary space is present in most forelimb bones in both *S. aegyptiacus* and *S. tenerensis* (***Figure 1****D, H*). The centra of anterior caudal vertebrae are occupied by a large medullary space (***Figure 1****C, J*), and large air-filled pneumatic spaces are present in the centra and neural arches of cervical vertebrae (***Evers et al. (2015***); ***Figure 1****B*). Collectively, these less dense, internal marrow- and air-filled spaces in *S. aegyptiacus* more than offset the added mass of infilled medullary space in the relatively reduced hind limb long bones (***Figure 1****E*). Hind limb bone infilling is better explained as compensation for the reduced size of the hind limb long bones that must support a body mass at the upper end of the range for theropods. Bending strength increases by as much as 35% when the medullary cavity is infilled (see ***Appendix 1***).

### *S. aegyptiacus* flesh model form and function

We added flesh to the skeletal model and divided the flesh model into body partitions adjusted for density. Muscle volume was guided by CT cross-sections from extant lizards, crocodylians and birds (***Figure 2****B*), and internal air space (pharynx-trachea, lungs, paraxial air sacs) was modeled on lizard, crocodilian and avian conditions (***Figure 2****C*; see Materials and Methods, ***Appendix 2***). Whole-body and body part surface area and volume were calculated, and body partitions were assigned density comparable to that in extant analogs (see Materials and Methods). For biomechanical analysis, we positioned the integrated flesh model in bipedal stance (***Figure 1****A*) as well as hybrid- and axial-powered swimming poses (***Grigg and Kirshner (2015***); ***Figure 2****A, B*).

The *center of mass* (CM) and *center of buoyancy* (CB) of the flesh model were determined to evaluate habitual stance on land and in shallow water (***Figure 1****A*), the water depth at the point of flotation (***Figure 2****D*), and its swimming velocity, stability, maneuverability, and diving potential in deeper water (***Figure 3***). No matter the included volume of internal air space, CM is positioned over the ground contact of symmetrically-positioned hind feet (***Figure 1****A*, red cross). Thus, *S. aegyptiacus* had a bipedal stance on land as previously suggested (***Henderson, 2018***), contrary to the aquatic hypothesis (***Ibrahim et al., 2020a***). Consistent with a bipedal stance, the manus is adapted for prey capture and manipulation (elongate hollow phalanges, scythe-shaped unguals) rather than weight support (***Figure 1****A, D*).

Adult *S. aegyptiacus* can feed while standing in water with flotation occurring in water deeper than ∼2.6 m (***Figure 2****D*). In hybrid or axial swimming poses, trunk air space tilts the anterior end of the model upward (***Figure 2****A, B*). With density-adjusted body partitions and avian-like internal air space, the flesh model of *S. aegyptiacus* has a body mass of 7,390 kg and an average density of 833 kg/m^3^ (see Materials and Methods), which is considerably less than the density of freshwater (1,000 kg/m^3^) and saltwater (1,026 kg/m^3^) or the average density of living crocodylians (1080 kg/m^3^; ***Grigg and Kirshner (2015***)).

*Swimming velocity* at the surface and underwater in extant lizards and crocodylians is powered by foot paddling and axial undulation (hybrid swimming) and at moderate to maximum (critical) speeds by axial undulation alone (axial swimming) (***Fish, 1984***; ***Grigg and Kirshner, 2015***). We used Lighthill’s (***1969***) bulk momentum formula to estimate maximum surface and underwater swimming velocity for the flesh model of *S. aegyptiacus*. Assuming a fully compliant *Alligator*-like tail (tail amplitude: 0.24/body length, tail wavelength: 0.57/body length and tailbeat frequency: 0.25 Hz; ***Fish (1984***); ***Sato et al. (2007***)), tail thrust (*P*_*t*_) and maximum velocity (*U*) can be determined:

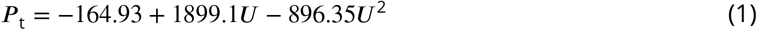

A body drag coefficient of 0.0035 assuming turbulent conditions was estimated for a Reynolds number of 752,400 at a swimming speed of 1.0 m/s. The total power from estimates of drag increased three-to-five fold to account for undulation of the tail, near-surface wave formation and increased sail drag when underwater (***Figure 3****A*). The addition of the sail increases the drag on the body of *S. aegyptiacus* by 33.4%. The intersection of the thrust power curve and drag power curves, where the animal would be swimming at a constant velocity, indicates slow maximum velocity at the surface (∼0.8 m/s) and only slightly greater when submerged (∼1.4 m/s) (***Figure 3****A*). Maximum tail thrust in S. aegyptiacus is 820 watts (683 N or 154 lbs), a relatively low value for the considerable caudal muscle mass in this large theropod (***Snivel and Russell, 2007***). Only a minor amount of caudal muscle power, however, is imparted to the water as thrust during undulation. As a result, maximum velocity is only 1.2 m/s, an order of magnitude less than extant large-bodied (>1m) pursuit predators. These species (mackerel sharks, billfish, dolphins and killer whales) are capable of maximum velocities of 10 to 33 m/s (***Tinsley, 1964***; ***Fish, 1998***; ***Fish and Rohr, 1999***; ***Iosilevskii and Weihs, 2008***).

*Stability* and the capacity to right are important in water. When positioned upright in water, the trunk sail of *S. aegyptiacus* is emergent (***Figure 3****B*, position 1). The flesh model, however, is particularly susceptible to long-axis rotation given the proximity of CM and CB, with stable equilibrium attained when floating on its side (***Figure 3****B*, position 3). Righting requires substantial torque (∼5,000 Nm) that is impossible to generate with vertical limbs and a tail with far less maximum force output (∼700 N). This stability predicament remains even with the smallest internal air space. The absence of vertical stability and righting potential in water stands in stark contrast to the condition in extant crocodylians and marine mammals (***Fish, 1998***; ***Grigg and Kirshner, 2015***).

*Maneuverability* in water (acceleration, turning radius and speed) wanes as body length increases (***Domenici, 2001***; ***Parson et al., 2011***; ***Domenici et al., 2014***; ***Hirt et al., 2017***; ***Gutarra and Rahman, 2022***), which is further compromised in *S. aegyptiacus* by its rigid trunk and expansive, unretractable sail. In contrast, large-bodied secondary swimmers capable of pursuit predation in open water have fusiform body forms with a narrow caudal peduncle for efficient tail propulsion (ichthyosaurs, cetaceans), control surfaces for reorientation, and narrow extensions (bills) to enhance velocity in close encounters with smaller more maneuverable prey (***Maresh et al., 2004***; ***Domenici et al., 2014***). Besides some waterbirds, semiaquatic pursuit predators are rare and include only the small-bodied (<2 m), exceptionally maneuverable otters that employ undulatory swimming (***Fish, 1994***).

*Diving* with an incompressible trunk requires a propulsive force (*F*_*p*_) greater than buoyancy. For *S. aegyptiacus*, in addition, a depth of ∼10 m is needed to avoid wave drag (***Figure 3****A*, bottom). The propulsive force required to dive is approximately 17,000 N:

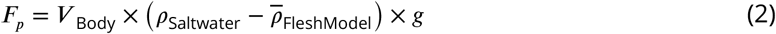

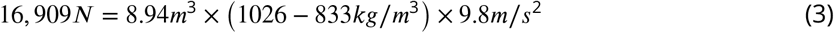

or approximately 25 times the maximum force output of the tail (*ρ*, density; 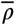, average density; *g*, gravitational acceleration). Even with lizard-like internal air space, diving still requires approximately 15 times maximum force output of the tail. To initiate a dive, furthermore, the tail would be lifted into the air as the body rotates about CB (***Figure 2****D*), reducing significant tail thrust. The now common depictions of *S. aegyptiacus* as a diving underwater pursuit predator contradict a range of physical parameters and calculations, which collectively characterize this dinosaur as a slow, unstable, and awkward surface swimmer incapable of submergence.

### Axial comparisons to aquatic vertebrates and sail-backed reptiles

Axial flexibility is requisite for axial-propulsion in primary or secondary swimmers. However, in *S. aegyptiacus*, trunk and sacral vertebrae are immobilized by interlocking articulations (hyposphenehypantrum), an expansive rigid dorsal sail composed of closely spaced neural spines and fused sacral centra (***Figure 1****A*).

The caudal neural spines in *S. aegyptiacus* stiffen a bone-supported tail sail by an echelon of neural spines that cross several vertebral segments (***Figure 4****A*). The caudal centra in *S. aegyptiacus* have nearly uniform subquadrate proportions along the majority of the tail in lateral view. These salient features of the tail suggest that it functioned more as a pliant billboard than flexible fluke.

No primary or secondary vertebrate swimmer has a comparable drag-magnifying, rigid dorsal sail, including sailfish, the dorsal fin of which is fully retractable and composed of pliable spines in membrane (***Domenici et al., 2014***). However, cranial crests and spine-supported torso-to-caudal sails have evolved multiple times among extant lizard (agamids, iguanians, chameleons) for intraspecific display rather than aquatic propulsion. Semiaquatic sailfin and basilisk lizards (***Figure 4****B*), for example, do not use their sails while swimming, spend very little time submerged, and are not aquatic pursuit predators.

In contrast, the distal tail of secondary swimmers such as crocodylians (***Grigg and Kirshner, 2015***), mosasaurs (***Lindgren et al., 2013***) and cetaceans (***Fish, 1998***) is expanded with pliable soft tissues free of bone to form a flexible caudal paddle or fluke (***Figure 4****C*). Likewise, caudal centra proportions in most secondary swimmers grade from subquadrate to spool-shaped in the distal half of the tail to increase flexibility and undulatory amplitude (***Figure 4****D*). Mosasaurs show a more derived piscine pattern of more uniform, disc-shaped centra (***Lindgren et al. (2013***); see ***Appendix 2***).

### Appendage comparisons to vertebrate secondary swimmers

Appendage surface area in secondarily aquatic axial swimmers is minimized to reduce drag, because terrestrial limbs are inefficient aquatic propulsors. Appendage surface area in *S. aegyptiacus*, in contrast, is substantially greater than in reptilian and mammalian secondary swimmers and even exceeds that of the terrestrial predators *Allosaurus* and *Tyrannosaurus* (***Figure 5***).

Interdigital webbing is used by some secondarily aquatic swimmers to increase the area of the foot paddle (***Fish, 2004***). Extant crocodylians use their limbs in paddling only at launch and slow speed before tucking them against the body (***Grigg and Kirshner, 2015***). Crocodylian interdigital webbing, which is better developed and always present in the hindfoot (***Figure 6****C, D*), only modestly increases surface area (<20%). Across a range of body size, we show that crocodylian paddle area scales isometrically (***Figure 6****F*; see ***Appendix 3***). Thus, the crocodylian foot paddle becomes even less effective as a propulsor with increasing body size. Nonetheless, a crocodylian of spinosaur size would have a foot paddle area an order of magnitude greater than is possible in *S. aegyptiacus* (***Figure 6****E*). Even a fully webbed hind foot in *S. aegyptiacus* (***Figure 6****A*) is far too small to have functioned as a significant aquatic propulsor or for stabilizing control.

### Paleohabitats and evolution

Most *Spinosaurus* fossils come from marginal basins along northern Africa in deltaic sediment laid down during an early Late Cretaceous transgression (***Figure 7****A*, sites 1, 2). These deposits, however, also include the majority of non-spinosaurid dinosaur remains, all of which may have been transported to some degree from inland habitats to coastal delta deposits. Because fossil transport is one way (downstream), documenting the inland fossil record is key to understanding true habitat range. We recently discovered fossils pertaining to *Spinosaurus* in two inland basins in Niger far from a marine coastline (***Figure 7***A, site 3). They were buried in fluvial overbank deposits alongside terrestrial herbivores (rebbachisaurid and titanosaurian sauropods) (see ***Appendix 4***).

The inland location of these fossils undermines the interpretation of *S. aegyptiacus* as a “highly specialized aquatic predator that pursued and caught its prey in the water column” (***Ibrahim et al., 2020a***). All large-bodied secondarily aquatic vertebrates — both extant (e.g., sea turtles, sirenians, seals, whales) and extinct (e.g., protostegid turtles, ichthyosaurs, metriorhynchoid crocodylomorphs, plesiosaurs) - are marine; none have been shown to live in both saltwater and freshwater habitats. Secondarily aquatic vertebrates that live in freshwater habitats (***Evers and Benson, 2019***; ***Motani and Vermeij, 2021***). Secondarily aquatic vertebrates that live in freshwater habitats have marine antecedents and are all small-bodied, such as river dolphins (<2.5 m length; ***Hamilton et al. (2001***)), small lakebound seals (<2 m; ***Fulton and Strobeck (2010***)), the riverbound Amazonian manatee (<2.5 m; ***Guterres-Pazin (2014***)), and a few mosasaurs and plesiosaurs of modest body size (***Gao et al., 2016***). In contrast, large-bodied semiaquatic reptiles frequent coastal and inland locales today and in the past. *Sarcosuchus imperator* is among the largest of semiaquatic reptiles (∼12 m length; ***Sereno et al. (2001***)) and lived in the same inland basin as *S. tenerensis*. The fossil record supports our interpretation of *Spinosaurus* as a semiaquatic bipedal ambush predator that frequented the margins of both coastal and inland waterways.

The large body size of *S. aegyptiacus* and antecedent species such as *S. tenerensis* also mitigates against an aquatic interpretation for the former, as it would constitute the only instance among vertebrates where the evolution of a secondarily aquatic species occurred at body size greater than 2-3 m. All other large-bodied secondarily aquatic vertebrates (e.g., ichthyosaurs, plesiosaurs, metriorhynchoid crocodylomorphs, protostegid turtles, mosasaurs, sirenians, whales) transitioned to an aquatic lifestyle at small body size, radiating subsequently within the marine realm to larger body size (***Domning, 2000***; ***Polcyn et al., 2014***; ***Moon and Stubbs, 2020***; ***Motani and Vermeij, 2021***; ***Thewissen et al., 2009***).

Phylogenetic analysis of an enlarged dataset for spinosaurids clarifies piscivorous adaptations in the earliest spinosaurids (stage 1) that enhance prey capture in shallow water and heighten visual display (***Figure 8***; ***Appendix 5***). In the skull, these include an elongate snout tipped with a dental rosette for snaring fish, retracted external nares to inhibit water intake, and a prominent nasal crest (***Charig and Milner, 1997***; ***Sereno et al., 1998***). The ornamental crest over the snout is accompanied by the evolution of a postcranial sail of varying height supported by neural spines of the posterior dorsals, sacrals and caudal vertebrae (***Stromer, 1915***; ***Sereno et al., 1998***; ***Allain et al., 2012***; ***Barker et al., 2021***). The earliest spinosaurids, in addition, have “cervicalized” anterior trunk vertebrae to enhance ventroflexion and the effective length of the neck, presumably as an adaptation to feeding in water (***Hone and Holtz, 2021***). Using the second dorsal vertebrae of the terrestrial predator *Allosaurus* for comparison, the homologous vertebra in spinosaurids shows marked modification (anterior face is convex, prominent ventral keel for muscular attachment, neural spine is reduced, zygapophyses large and planar). Giraffids, for a similar purpose, have “cervicalized” the first thoracic vertebra to facilitate dorsiflexion and effective neck length (***Lankester, 1908***; ***Danowitz et al., 2015***; ***Müller et al., 2021***).

Other features present in stage 1 indicate that baryonychines (e.g., *Suchomimus, Baryonyx, Ceratosuchops*) have a low nasal crest and swollen brow ridges as ornamentation on the skull for display or agonistic purposes, respectively. Neural spines over the trunk and tail are heightened to a varying degree (***Figure 1****B*). Spinosaurines (***Figure 1****A*; ***Figure 8***, stage 2) exhibit further specializations for display (heightened cranial crest, low cervical sail, hypertrophied torso-to-caudal sail) and piscivory (spaced teeth with smooth carinae, smaller, more retracted external nares, scythe-shaped manual unguals.

## Conclusions

1. Adult *S. aegyptiacus* had a body length of approximately 14 m with the axial column in neutral pose.
2. The reduced hind limb long bones in *S. aegyptiacus* are infilled to a varying degree probably as an adaptation to weight support on land, because the added mass from infilling is more than offset by significant paraxial pneumaticity along the length of the presacral column. Medullary cavities, in addition, are common in forelimb long bones and in anterior caudal centra.
3. The segment-crossing caudal neural spines in *S. aegyptiacus* suggest that its tail functioned more as a pliant billboard than flexible fluke. Similar spine-supported torso-to-caudal sails in extant reptiles are used for display rather than in swimming. The expanded keels on the tail of extant and extinct vertebrate secondary swimmers, in contrast, is composed mostly of pliable soft tissues free of bone.
4. *S. aegyptiacus*, like *S. tenerensis* and other spinosaurids, was bipedal on land with its center of mass positioned over its hind feet. The long-clawed forelimbs of *S. aegyptiacus* were not used in weight support on land.
5. An adult flesh model of *S. aegyptiacus* has a body mass of ∼7400 kg and average density of ∼830 kg/m^3^, which is considerably less than the density of saltwater (1,026 kg/m^3^).
6. *S. aegyptiacus* could wade into shallow water for feeding with flotation occurring at water depth greater than ∼2.6 m.
7. *S. aegyptiacus* was incapable of diving, given its buoyancy and incompressible trunk. Full sub-mergence would require 15-25 times the maximum force output of its tail, depending on estimated lung volume.
8. *S. aegyptiacus* was unstable in deeper water with little ability to right itself, swim, or maneuver underwater. Maximum power from its tail, assuming it could undulate as in *Alligator*, is less than 700N, which would generate a top speed of ∼1m/s, an order of magnitude slower than extant large-bodied pursuit predators.
9. All extant and extinct large-bodied (>2m long) secondarily aquatic vertebrates are strictly marine, whereas fossils pertaining to *Spinosaurus* have been found in inland basins distant from a marine coast.
10. Transition to a semiaquatic lifestyle, as occurred in the evolution of spinosaurid theropods, can occur at any body size. In contrast, transition to an aquatic lifestyle among tetrapods has only occurred at relatively small body size (<3 m) with subsequent radiation once in the marine realm into larger body sizes.
11. *S. aegyptiacus* is interpreted as a semiaquatic shoreline ambush predator more closely tied to waterways than baryonychine spinosaurids.
12. Spinosaurids flourished over a relatively brief Cretaceous interval (∼35 My) in circum-Tethyan habitats with minimal impact on aquatic habitats globally.
13. Two phases are apparent in evolution of aquatic adaptations among spinosaurids, the second distinguishing spinosaurines as the most semiaquatic (but not aquatic) non-avian dinosaurs.

## Materials and Methods

### Skeletal reconstruction

The composite skeletal reconstruction of *S. aegyptiacus* is based principally on bones of holotypic and neotypic specimens supplemented by associated and isolated bones from Cenomanian-age formations in Egypt, Morocco and Niger (***Figure 1****A*). The two most important specimens include the subadult partial skeleton composing the holotype (BSP 9012 VIII 19) from the Western Desert of Egypt (***Stromer, 1915***; ***Smith et al., 2006***) and a subadult partial skeleton designated as the neotype from the Kem Kem Group in Morocco (FSAC-KK 11888; ***Ibrahim et al. (2014***, 2020b)). A third referred specimen from Egypt was also considered (BSPG 1922 X45, “*Spinosaurus* B”; ***Stromer (1934***). These are the only associated specimens known for *Spinosaurus aegyptiacus* on which to base the skeletal reconstruction, the relative size calculated from overlapping bones (***Table 1***). Of these three specimens, only the bones of the neotype are preserved, all of which have been CT-scanned except for recently discovered bones of the tail (***Ibrahim et al., 2020a***). Noteworthy isolated specimens have been recovered from the Kem Kem Group in Morocco, including a large snout and manual phalanx used to gauge maximum adult body size.

**Table 1.** Relative size of specimens in the skeletal reconstruction of *S. aegyptiacus*. Relative sizes of key specimens used in the skeletal model of *S. aegyptiacus* (nos. 1-4) and select bones (nos. 5, 6) from Egypt and Morocco. All are scaled to the size of the adult snout (MSMN V4047).

| No. | Specimen | Maturity | Relative Size | Linear Up-sizing | Description |
| --- | --- | --- | --- | --- | --- |
| 1 | BSPG 1912 VIII 19 | subadult | 76% | 1.32 | Holotype (destroyed) preserving dentaries, presacral, sacral and caudal vertebrae including the dorsal sail (*Stromer, 1915*) |
| 2 | FSAC-KK 11888 | subadult | 76% | 1.32 | Neotype preserving skull bones, partial limbs, dorsal sail and most of the tail (*Ibrahim et al., 2014*) |
| 3 | BSPG 1922 X 45 | subadult | 66% | 1.51 | "*Spinosaurus* B" (destroyed) fragmentary specimen with 5 partial dorsals (~D1 centrum, mid dorsal centrum, partial ~D13 vertebra), 7 partial caudal vertebrae, and both tibiae (*Stromer, 1934*) |
| 4 | MSMN V4047 | adult | 100% | — | Isolated snout with broken teeth (*Dal Sasso et al., 2005*) |
| 5 | UCRC PV8 | adult | 80% | 1.25 | Large manual phalanx I-1 (28.0 cm length) of an adult within reach but still smaller than the 35 cm length estimated on the basis of the proportions in the manus of *Angaturama* (? =*Irritator*) scaled to the adult snout (*Aureliano et al., 2018*) |
| 6 | UCRC PV24 | adult | 75% | 1.33 | Large Kem Kem vertebra, ~C9, from Gara Sbaa (centrum length 11.6 cm, centrum width 14.0 cm) |

We incorporated all CT-scanned bones of the neotype and reconstructions (based on lithographic plates and photographs) of bones of the holotype and referred specimen from Egypt. For unknown bones without sequential adjacency as a guide, other spinosaurids were consulted for shape and proportion. In the case of overlapping bones, priority was given to the neotype (blue) followed by the holotype (red) and referred specimens (yellow). Bones without representation among specimens attributed to *Spinosaurus aegyptiacus* are shaded gray (***Figure 1****A*).

### Skeletal reconstructions compared

We compared our digital skeletal model of *S. aegyptiacus* to the recently published two-dimensional silhouette skeletal reconstruction in the aquatic hypothesis (***Ibrahim et al., 2020a***), both of which are based primarily on holotypic and neotypic specimens (***Figure 9***). We registered the reconstructions to each other by superimposing the four longest complete bones of the neotype (femur, tibia, ilium, ischium). Significant differences are apparent in several dimensions with major implications for the calculation of center of mass and buoyancy.

**Figure 9.**
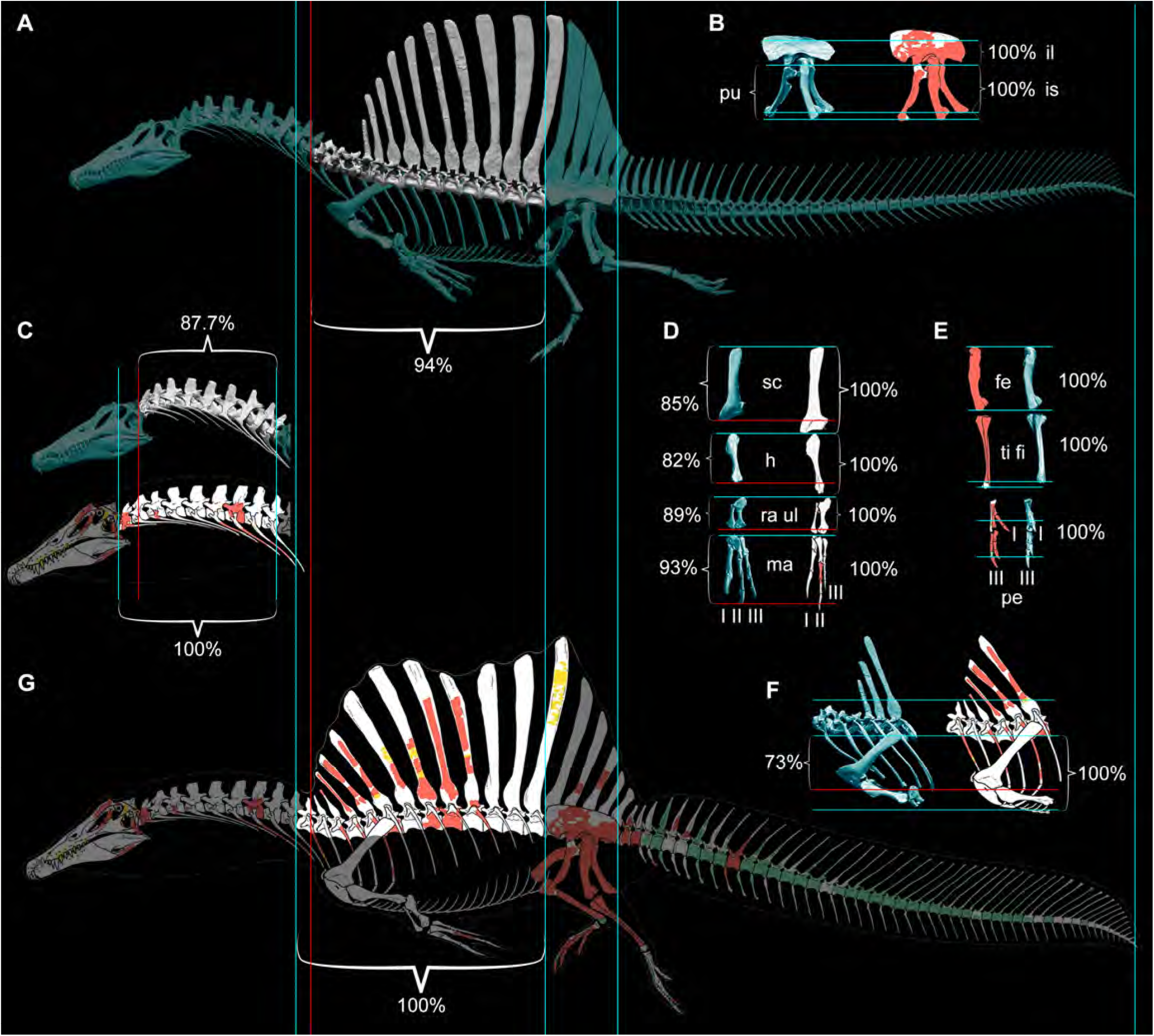
Comparison of skeletal reconstructions for *S. aegyptiacus* in left lateral view. **(A)** Digital skeletal reconstruction from this study in left lateral view. **(B)** Pelvic girdle. **(C)** Cervical column (C1-10). **(D)** Pectoral girdle and forelimb. **(E)** Hind limb. **(F)** Anterior trunk. **(G)** Silhouette skeletal drawing from the aquatic hypothesis (from ***Ibrahim et al. (2020a***)). On one side of each length comparison, one or two blue lines are shown that register the alternative reconstructions. The opposing end of each length comparison either has a single blue line (when comparisons match, both 100%) or a red line as well for the shorter one (<100%). Skeletal reconstructions (A,G) are aligned by the anterior and posterior margins of the ilium and measured to the cervicodorsal junction (C10-D1); the pelvic girdle (B) is aligned along the ventral edge of the sacral centra and base of the neural spines and measured to the distal ends of the pubis and ischium; the cervical column (C) is aligned at the cervicodorsal junction (C10-D1) and measured to the anterior end of the axis (C2); the scapula and components of the forelimb (humerus, ulna, manual digit II, manual phalanx II-1) (D) are aligned at the distal end of the blade and their proximal ends, respectively, and measured to the opposing end of the bone; the components of the hind limb (femur, tibia, pedal digits I, III) (E) are aligned at their proximal ends and measured to the opposing end of the bone; and anterior trunk depth (F) is aligned along the ventral edge of the centrum and neck of the spine of D6 and measured to the ventral edge of the coracoid. All limb bones compared are from the left side with the exception of the pes, which is from the better preserved right side of the neotype. Abbreviations: I, II, III, digits I-III; II-1, phalanx II-1; fe, femur; h, humerus; il, ilium; is, ischium; ma, manus; pe, pes; pu, pubis; sc, scapula; ti, tibia; ul, ulna.

When aligned at the hip, sacral and caudal columns have nearly identical length, but the presacral column is significantly longer (∼10%) in the reconstruction of the aquatic hypothesis. The extra length of the presacral column is located in the neck between C2-10 and torso between D4-13. The trunk in our digital skeletal model is also not as deep as that in skeletal silhouette drawing, as can be seen by aligning the skeletons along the dorsal column (***Figure 9****C*). The contour of the belly marked by the gastral basket and the coracoids of the pectoral girdle extend farther ventrally (∼25%) than the ends of the pubes, unlike our digital reconstruction or that of most other silhouette reconstructions for non-avian theropods. The length of the ribcage in our model are consistent with the only well preserved spinosaurid ribcage known to date (*Suchomimus tenerensis*, MNBH GAD70).

Finally, the forelimb in the skeletal silhouette drawing is approximately 30% longer than that in our digital reconstruction (***Figure 9****A*). The neotype is the only associated specimen of *S. aegyptiacus* preserving bones from the forelimb (partial manual digit II). The preserved manual phalanges are slender with deeply cleft distal condyles, which allows reference of additional phalanges of similar form from the Kem Kem Group (***Figure 1****D*). Our reconstruction of the manus is based on a recently described forelimb of the close relative *Irritator* (= *Angaturama*; ***Machado and Kellner (2009***); ***Aureliano et al. (2018***)). The proportions of more proximal forelimb segments and the pectoral girdle are based on the holotypic specimens of *Baryonyx* and *Suchomimus*. The forelimb in *S. aegyptiacus* is robust and long relative to other non-avian theropods, although considerably shorter than in some previous reconstructions (***Ibrahim et al., 2014, 2020a***).

The longer presacral proportions, deeper torso, and longer forelimb of the skeletal reconstruction and flesh model used by the aquatic hypothesis cantilever significant additional body mass anterior to the hip joint. That additional front-loading appears to be the main factor generating their mid trunk location for center of mass and the basis for regarding *S. aegyptiacus* as quadrupedal on land (***Ibrahim et al., 2020a***).

### Flesh reconstruction of axial musculature

To estimate the volume of axial musculature in *S. aegyptiacus* (***Figure 10***), we referenced CT-based studies on the ostrich (*Struthio*; ***Wedel (2003***); ***Snively and Russell (2007***); ***Persons IV et al. (2020***)) and alligator *Alligator* (***Cong, 1998***; ***Mallison et al., 2015***). To estimate caudal muscle mass, we used CT scans of various reptiles including the sail-backed basilisk lizard, *Basilicus plumifrons* (***Figure 11, Table 2-5***).

**Table 2.**
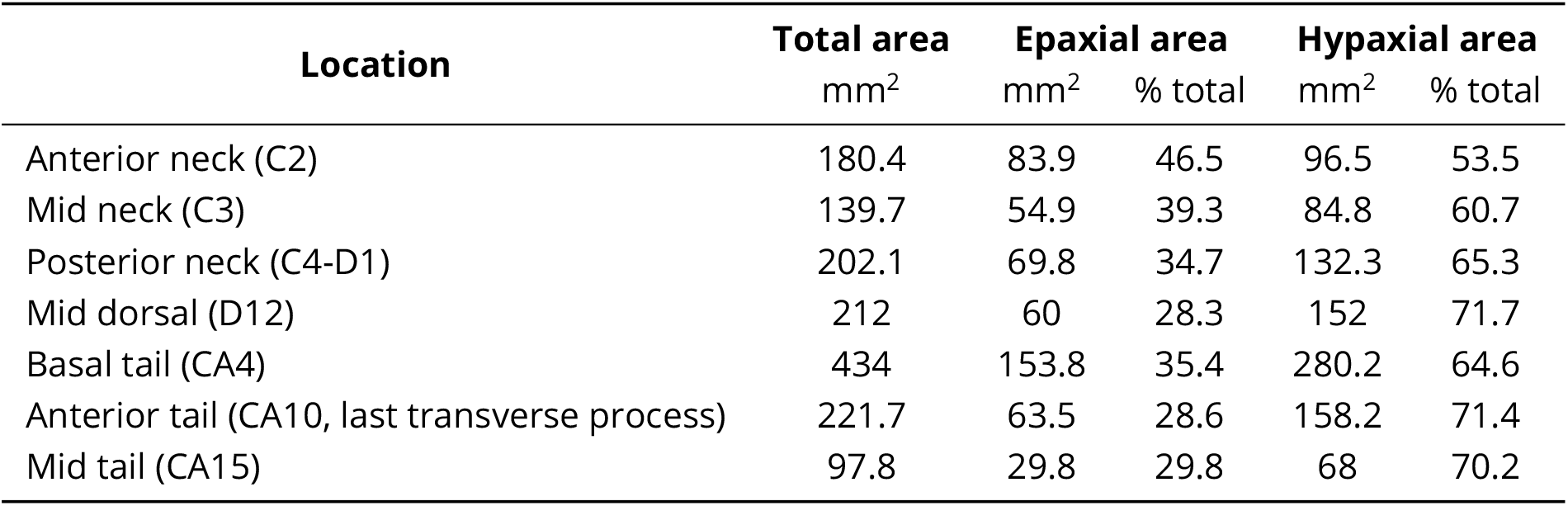
Axial muscle area in the crested basilisk. Area measurements of epaxial and hypaxial musculature along the axial column in the crested basilisk *Basiliscus plumifrons* (FMNH 112993). C, cervical; D, dorsal; CA, caudal.

**Table 3.** Axial muscle and transverse process length in the crested basilisk. Transverse processes versus muscle width in the tail cross-sections in the crested basilisk *Basiliscus plumifrons* (FMNH 112993). Measurements are from the midline to the distal end of the transverse process (or centrum margin when there is no process) and to the lateral surface of the tail.

| Location | Measurement | Total (mm) | % bone and muscle | % muscle only |
| --- | --- | --- | --- | --- |
| CA4 | transverse process width | 16.8 | 62.9 | 37.1 |
|  | total width | 26.7 |  |  |
| CA10 | transverse process width | 8.1 | 57 | 43 |
|  | total width | 14.2 |  |  |
| CA15 | centrum width | 3.5 | 36.8 | 63.2 |
|  | total width | 9.5 |  |  |

**Table 4.**
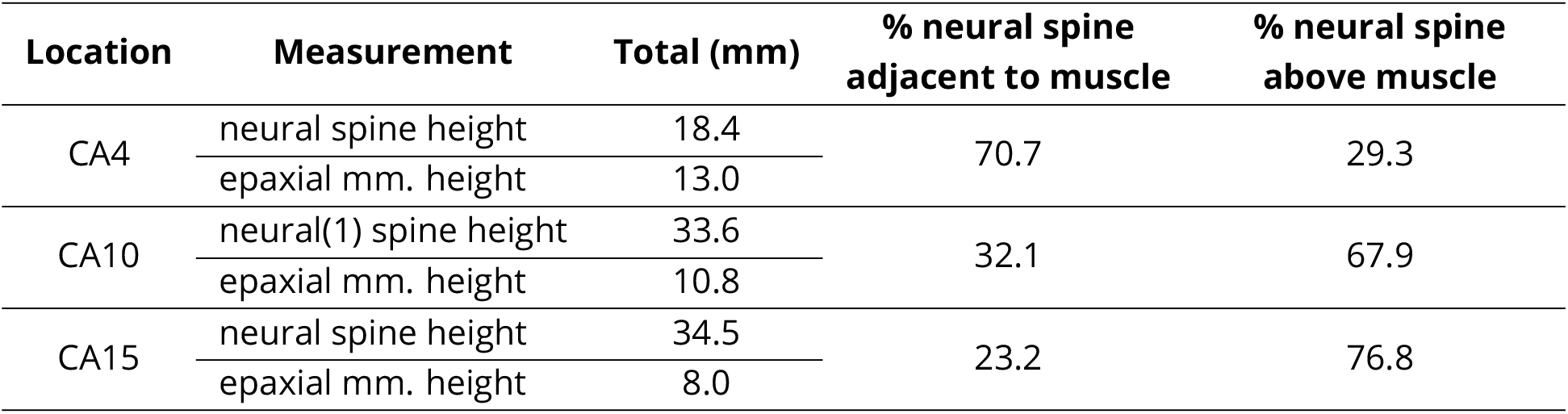
Epaxial muscle height and neural spine height in the crested basilisk. Height of neural spines versus epaxial musculature in tail cross-sections in the crested basilisk *Basiliscus plumifrons* (FMNH 112993). Measurements are from the dorsal surface of the centrum to the top of the epaxial muscle and to the distal end of the neural spine.

| Location | Measurement | Total (mm) | % neural spine adjacent to muscle | % neural spine above muscle |
| --- | --- | --- | --- | --- |
| CA4 | neural spine height | 18.4 | 70.7 | 29.3 |
|  | epaxial mm. height | 13.0 |  |  |
| CA10 | neural(1) spine height | 33.6 | 32.1 | 67.9 |
|  | epaxial mm. height | 10.8 |  |  |
| CA15 | neural spine height | 34.5 | 23.2 | 76.8 |
|  | epaxial mm. height | 8.0 |  |  |

**Table 5.** Hypaxial muscle depth and chevron length in the crested basilisk. Chevron length versus hypaxial muscle depth in tail cross-sections in the crested basilisk *Basiliscus plumifrons* (FMNH 112993). Measurements are from the ventral surface of the centrum to the distal tip of the chevron and to the ventral surface of the tail.

| Location | Measurement | Total (mm) | % chevron length | % muscle below chevron |
| --- | --- | --- | --- | --- |
| CA4 | chevron depth | 9 | 52.6 | 47.4 |
|  | hypaxial depth | 17.1 |  |  |
| CA10 | chevron depth | 7.1 | 67 | 33 |
|  | hypaxial depth | 10.6 |  |  |
| CA15 | chevron depth | 4.8 | 67.6 | 32.4 |
|  | hypaxial depth | 7.1 |  |  |

**Figure 10.**
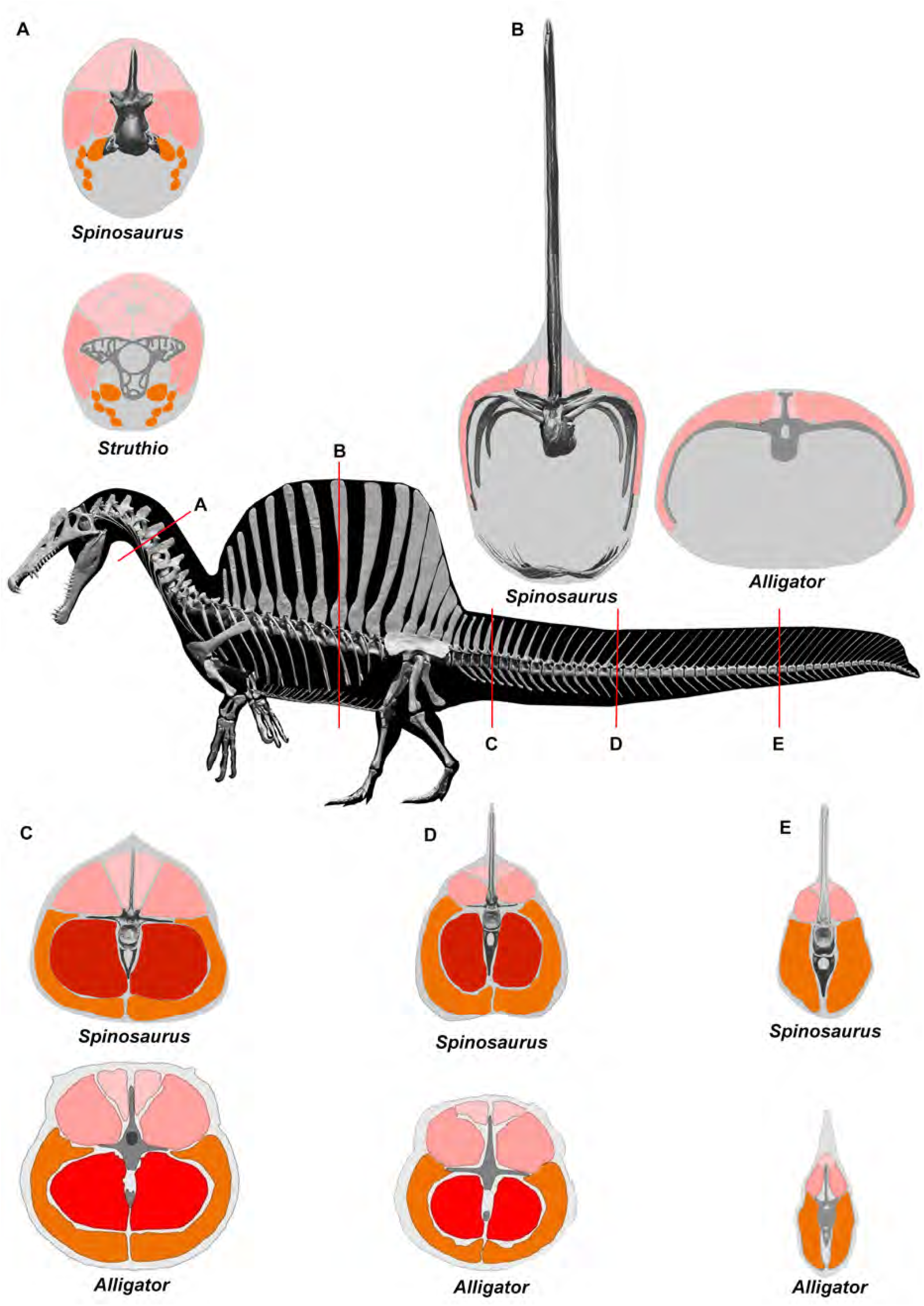
**Cross sections for the flesh model of** S. aegyptiacus **compared to CT-based cross sections from ostrich and alligator** (after ***Wedel (2003***); ***Böhmer et al. (2019***); ***Mallison et al. (2015***); ***Cong (1998***)). Cross sections at (A) Mid neck, (B) Mid trunk, (C) Anterior tail, (D) Mid tail, and (E) Distal tail. Epaxial muscles pink; hypaxial muscles orange; caudofemoralis muscle in red.

**Figure 11.**
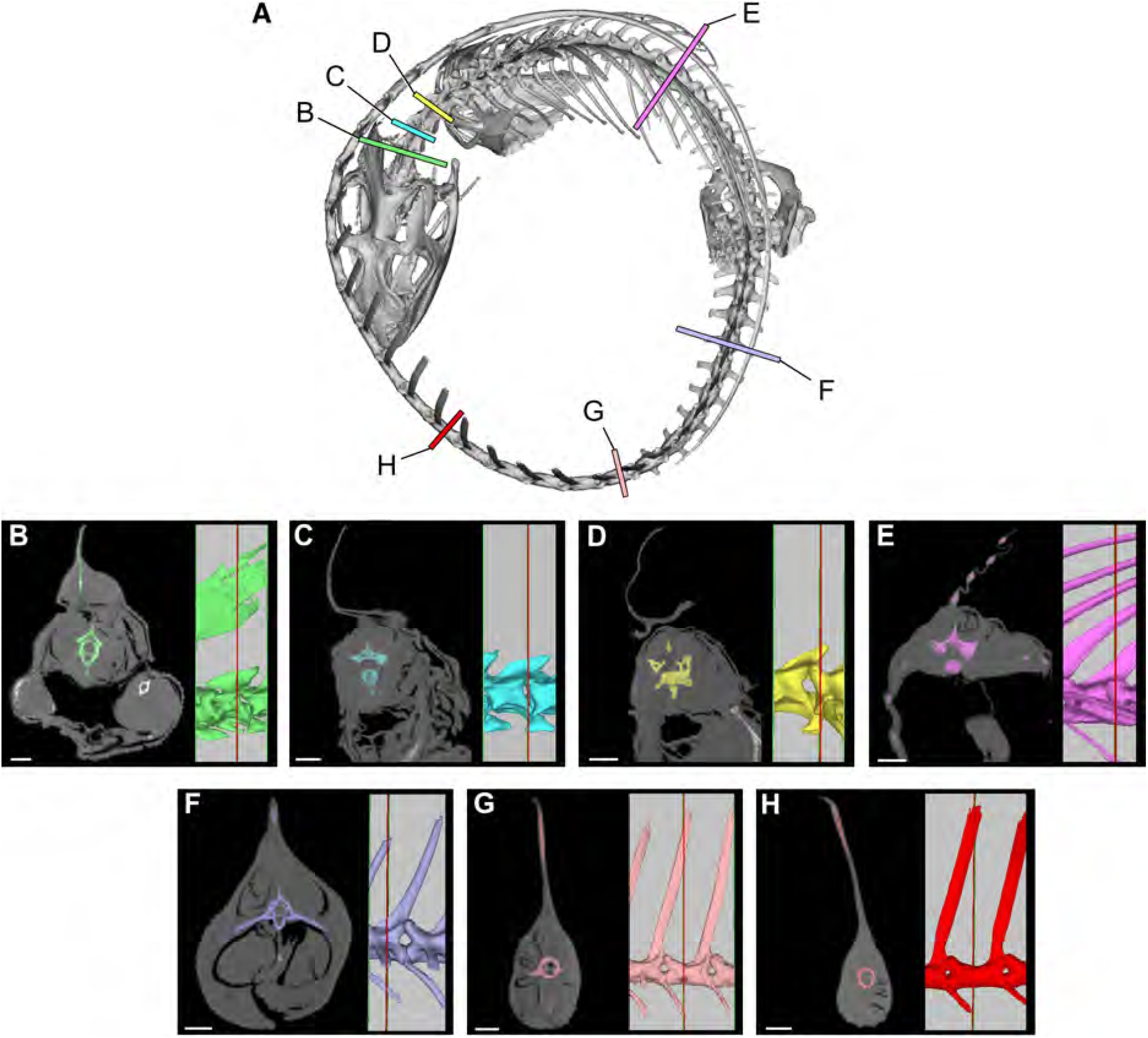
Cross-sections from a CT scan of a male *Basiliscus plumifrons* FMNH 112993. **(A)** Skeleton showing position of computed-tomographic sections of the axial column. **(B)** Anterior cervical region (C2). **(C)** Mid cervical region (C3). **(D)** Posterior cervical and anterior dorsal region (C4-D1). **(E)** Mid dorsal region (D12). **(F)** Anterior caudal region (CA4). **(G)** Caudal region at most posterior transverse process (CA10). **(H)** Mid caudal region (CA15). CT scan data available on Morphosource.org. Abbreviations: C, cervical; CA, caudal; D, dorsal. Scale bars, 5mm.

For epaxial muscle mass in *S. aegyptiacus*, we estimated its vertical extent as twice centrum height, measuring upward from the base of the neural spine. The transverse width of epaxial musculature was estimated to be a little less than that of the hypaxial muscles, widest ventrally and tapering to the midline dorsally. For hypaxial muscle mass in *S. aegyptiacus*, we estimated its vertical depth at approximately twice chevron length in the anterior tail and 1.5 times chevron length in mid and posterior portions of the tail. We estimate the transverse width of hypaxial muscles as twice the length of the transverse processes. CT cross-sections of extant reptiles show that considerable muscle mass is present beyond the distal end of caudal transverse processes in anterior and middle portions of the tail (***Figure 10***).

Several cross-sections from *Basiliscus plumifrons* (crested basilisk) provided valuable insights on the distribution of axial muscles in a lizard with a dorsal-to-caudal sail (***Figure 11, Table 2-5***). Epaxial musculature in the trunk and tail comprises less than one-third of total axial muscle volume (***Table 2***). Caudal neural spines project beyond the epaxial musculature to support the sail to a greater extent in mid and distal portions of the tail. At the base of the tail (CA4), approximately one-third (29.3%) of the neural spine projects dorsally supporting the sail. At mid tail (CA15), approximately three-quarters (76.8%) of the neural spine projects dorsally supporting the sail. The hypaxial musculature extends well below the distal end of the chevrons. At the base of the tail (CA4), the chevron lies internal to approximately one-half (52.6%) of hypaxial muscle depth, with the remainder (47.4%) distal to the end of the chevron. Farther along the tail (CA10-15), the chevrons are proportionately longer, supporting approximately two-thirds (67%) of hypaxial muscle depth with approximately one-third of hypaxial musculature beyond the distal end of the chevron. These cross-sections confirm the presence of considerable muscle mass ventral to the distal end of the chevrons in anterior through mid portions of the tail (***Figure 10, Figure 11***).

### Flesh model density partitions, dimensions and properties

The digital skeletal model was wrapped in flesh (using ZBrush) guided by recent documentation of muscle mass in CT scans of various modern analogs (***Wedel, 2003***; ***Snively and Russell, 2007***; ***Mallison et al., 2015***; ***Persons IV et al., 2020***; ***Díez Díaz et al., 2020***). We inserted anatomically-shaped and -positioned air spaces (pharynx-trachea, paraxial air sacs, lungs) of optional volumes (minimum-lizard, medium-crocodylian, maximum-avian) within the head, neck and torso (***Figure 2****C*). For additional measurements, we added a “mesh” over the flesh model (***Henderson (1999***); ***Figure 2****D*).

We divided the flesh model into six parts (axial body-presacral, axial-caudal, dorsal-to-caudal sail, forelimbs, hindlimbs, lungs) in order to assign appropriate densities (***Table 6***). Densities were assigned to body parts based on their estimated composition, using values for tissues ranging from fat (900 gm/l) to compact bone (2,000 gm/l). The average whole body density for *S. aegyptiacus*, 833 gm/l (***Table 6***), compares favorably to whole body density estimates for various non-avian dinosaurs (800-900 gm/l). We have compiled various functional dimensions of the adult flesh model (***Table 7***), and we divided the flesh model into 10 body parts, for which we list volumes and external surface areas (excluding cut surfaces) (***Table 8***).

**Table 6.** Density, volume and mass in the flesh model of *S. aegyptiacus*. Whole body and body part densities, volumes and masses for the new mesh adult flesh model of *S. aegyptiacus*.

| No. | Body Partition | Average density (kg/m³) | % of Axial volume without sail | Mass (kg) |
| --- | --- | --- | --- | --- |
| 1 | Whole body | 833 | — | 7,390 |
| 2 | Axial body (excluding lung/sail) | 788 | 100 | 5,794 |
| 3 | Axial head-trunk (not lung/sail/tail) | 850 | 64.8 | 3,209 |
| 4 | Axial tail (not sail) | 1,000 | 35.2 | 2,585 |
| 5 | Forelimb (paired) | 1,050 | 3.8 | 108 |
| 6 | Hind limb (paired) | 1,050 | 11.1 | 590 |
| 7 | Dorsocaudal sail | 1,196 | 8.5 | 441 |
| 8 | Lungs | 0 | 12.5 | 0 |

**Table 7.**
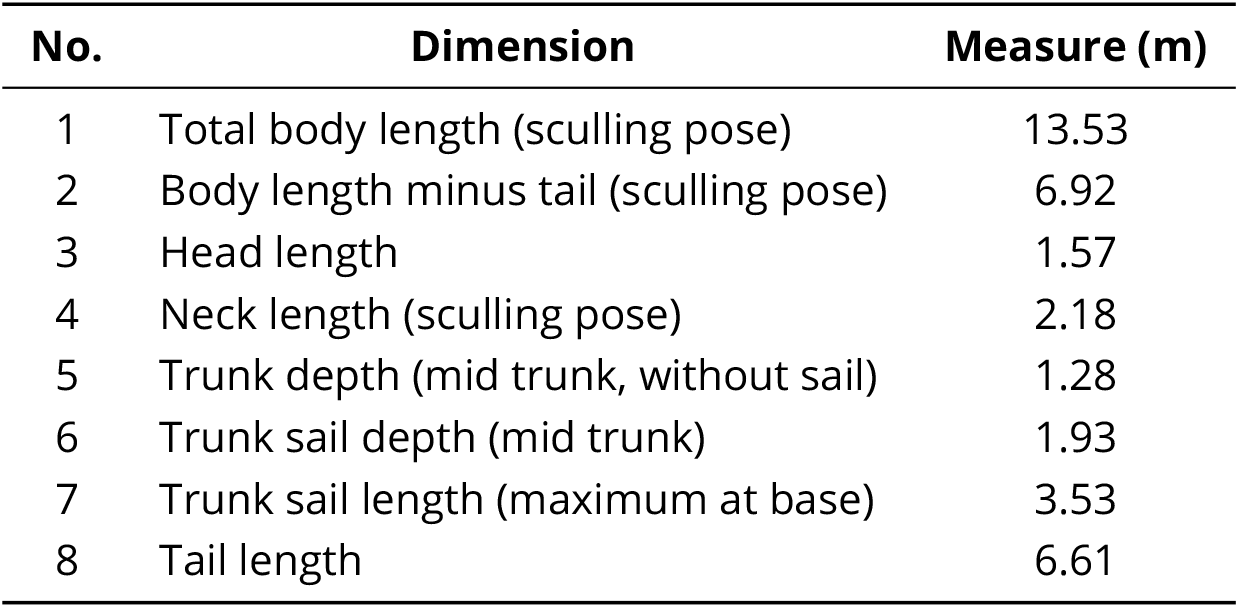
Flesh model functional dimensions in *S. aegyptiacus*. Functional dimensions for the adult flesh model of *S. aegyptiacus* in sculling pose.

| No. | Dimension | Measure (m) |
| --- | --- | --- |
| 1 | Total body length (sculling pose) | 13.53 |
| 2 | Body length minus tail (sculling pose) | 6.92 |
| 3 | Head length | 1.57 |
| 4 | Neck length (sculling pose) | 2.18 |
| 5 | Trunk depth (mid trunk, without sail) | 1.28 |
| 6 | Trunk sail depth (mid trunk) | 1.93 |
| 7 | Trunk sail length (maximum at base) | 3.53 |
| 8 | Tail length | 6.61 |

**Table 8.** Flesh model volume and surface area in *S. aegyptiacus*. Adult flesh model whole body and body part volumes and surface areas as measured in MeshLab. Surface area of body parts does not include cut surfaces.

| No. | Body Part | Volume (m<sup>3</sup>) | Surface Area (m<sup>2</sup>) |
| --- | --- | --- | --- |
| 1 | Whole body | 8.94 | 54.06 |
| 2 | Body above waterline (floating) | 1.65 | 22.58 |
| 3 | Body below waterline (floating) | 7.27 | 31.38 |
| 4 | Head | 0.21 | 2.23 |
| 5 | Neck | 0.78 | 4.37 |
| 6 | Trunk | 4.01 | 11.44 |
| 7 | Trunk sail (both sides, external edge) | 0.40 | 10.06 |
| 8 | Forelimb (both) | 0.24 | 3.86 |
| 9 | Hind limb (both) | 0.45 | 5.29 |
| 10 | Tail | 2.81 | 16.56 |
| 11 | Tail with axial muscle | 2.71 | 13.27 |
| 12 | Tail sail only | 0.10 | 3.17 |
| 13 | Airspace-minimum (~4% volume) | 0.37 | 4.86 |
| 14 | Airspace-medium (~8% body volume) | 0.67 | 6.63 |
| 15 | Airspace-maximum (~12% body volume) | 1.08 | 8.66 |

We registered center of mass (CM) as the horizontal distance from the apex of the acetabulum (*x*-coordinate) and the vertical distance from the ground surface under the sole of the foot (*y*-coordinate) (***Table 9***, no. 4). This is the fourth estimation of center of mass for *S. aegyptiacus*, and we argue here the most accurate. We prefer a measure from the acetabulum rather than the distal tail tip, which as in *S. aegyptiacus* is often a matter of speculation given the rarity of completely preserved caudal columns (***Hone, 2012***). For the acetabulum, we recommend using its “apex” rather than its “cranial end” (***Ibrahim et al., 2020a***) for three reasons. First, the apex of the acetabulum is a more easily recognized landmark than the poorly defined anterior edge (or rim) of the acetabulum. Second, the apex rather than the “cranial end” of the acetabulum, is a more functionally intuitive point from which to measure center of mass, given its proximity to the rotational point for body mass centered over the hind limbs. And third, the dorsal (proximal) articular end of the femoral head is close to the apex of the acetabulum, and so length the femur and the distance that CM lies farther forward can be directly compared (CM located anteriorly beyond femoral length excludes stable bipedal posture with a relatively horizontal dorsosacral column).

**Table 9.** Center of mass (CM) calculations for *S. aegyptiacus*. There have been four estimates for the location of CM in flesh models of *S. aegyptiacus* using four different points of origin as a reference. Because three were based on an adult flesh model, we convert the one study based on a subadult (number 3) to reflect its position in an adult flesh model.

| No. | Author/Result | x-origin | CM (cm)<br>(x = anterior to origin) |  | Notes |
| --- | --- | --- | --- | --- | --- |
|  |  |  | x | y |  |
| 1 | *Ibrahim et al. (2014); quadruped* | hip joint | >81 | — | Based on an adult flesh model, no coordinates given, CM shown graphically under D10 and said to be anterior to hip/knee joints at a distance greater than femur length (MeshLab calculation error) |
| 2 | *Henderson (2018); biped* | tip of tail | 8,850 | 100 | Based on a 3D mesh model based on the adult skeletal model of Ibrahim et al. (2014) with estimated length 16 m long; y-coordinate origin is “lowest point of axial body” |
| 3 | *Ibrahim et al. (2020a); quadruped* | “cranial rim” of acetabulum | 72.5 - 82.5<br>(adult = 95.7-108.9) | -81<br>(106.9) | Based on a flesh model of the subadult neotype with femur length of 62.5 cm (actual 61.0 cm); y-coordinate measures to substrate |
| 4 | this paper; biped | apex of the acetabulum | 15.3 | -240 | Based on an adult flesh model with avian-style internal air spaces and femur length of 81.0 cm; y-coordinate measures to substrate |

With avian-like air space (“maximum”), CM is positioned only 15.3 cm anterior to the apex of the acetabulum and clearly over the pedal phalanges of the foot for a bipedal stance (***Table 10***). The smallest air space option modeled on lizards (“minimum,” only 4% of body volume) generates the heaviest torso and displaces CM anteriorly 13.2 cm to a distance of 28.5 cm from the apex of the acetabulum (***Table 10***). In this location, CM is still approximately 12 cm short of the midpoint along the length of the femur (∼40 cm; femoral length is 81 cm in adult *S. aegyptiacus*). In this worst-case scenario regarding internal air volume, CM is still positioned over the pedal phalanges of the hind limb. Our flesh model does not support an obligatory quadrupedal pose on land for *S. aegyptiacus*.

**Table 10.** Estimated internal air space in *S. aegyptiacus*. Air space options for the adult flesh model of *S. aegyptiacus* and their effect on whole body density, body mass (BM), and center of mass (CM). The *x*-coordinate for CM is measured from the apex of the acetabulum.

| No. | Air space option | Part of whole-body volume (%) | Mean whole-body density (Gm/l) | BM (kg) | CM |
| --- | --- | --- | --- | --- | --- |
|  |  |  |  |  | x-coordinate (cm) |
| 1 | Minimum (lizard-like) | 4 | 909 | 8,013 | 28.5 |
| 2 | Medium (croc-like) | 8 | 875 | 7,716 | 23.2 |
| 3 | Maximum (bird-like) | 12.5 | 833 | 7,390 | 15.3 |

## Acknowledgments

We thank A Resetar, J Mata and D Coldren of the Field Museum of Natural History, R Carmichael of the Wildlife Center and R Bavrisha of the Chicago Herpetological Society for access to recent preserved and live reptile specimens; Morphosource, oVert TCN, UMMZ, KU, and UF for access to and archiving of CT scans; R Shonk and A Schulte for digital modeling; E Fitzgerald for fossil preparation; J Mallon for photography; and E Saitta, E Johnson-Ransom, DB Dutheil, SW Evers, E Snively, and J Schwartz for comments on the manuscript. This research was supported by Bob and Ellen Vladem and SC Johnson.

## Additional information

### Author contributions

Conceptualization, PCS, NM, DMH, FEF, DV, SLB and KKF; Initial draft, PCS; Review and editing, DMH, DV, SLB, FEF, KKF and NM; Biomechanical simulation, DMH, NM and FEF; Fossil CT imaging, skeletal and flesh model reconstruction, TMK, LLC and DV; Modern analog CT imaging and data collection, SLB, DV; Final figures, LLC and PCS with contributions from SLB, DMH, FEF, DV and NM; Phylogenetic analysis, DV and PCS.

### Decision letter and Author response

Decision letter

Author response

## Additional files

### Supplementary files

#### Data availability

All data generated or analyzed in this study are included in the paper, Appendices, and Morphosource.

## Appendix 1

### Long bone infilling in *Spinosaurus aegyptiacus*

In *S. aegyptiacus* the medullary cavities in the long bones of the hind limb are reduced in diameter or infilled altogether by non-pachystotic bone. This condition, which is unusual among non-avian dinosaurs, was initially thought to be an adaptation for decreasing buoyancy (***Ibrahim et al., 2014***; ***Fabbri et al., 2022***), although that function has been challenged recently (***Myhrvold et al., in press***). Enhanced bone strength is an alternative explanation for solid hind limb long bones in *S. aegyptiacus*, a bipedal theropod of very large body size with a reduced hind limb. Solid long bones are known to occur in other large-bodied bipedal and quadrupedal dinosaurs and mammals (***Vanderven et al., 2014***; ***Houssaye et al., 2016***).

To better understand this alternative explanation, we compare the increase in strength of a femur with solid shaft compared to one of identical size with a hollow shaft, using the section modulus as an indicator of the resistance to bending (***Farlow et al., 1995***). For a solid circular cross-section, the section modulus (*SM*) is given by 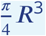, where *R* is the outer radius. When *R* has a value 1.0, *SM* is 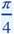, or 0.785 m^3^. The *SM* for a cylinder with a hollow core is calculated by subtracting the *SM* of an inner cylinder comprising the hollowed space from that of the outer cylinder.

An inner radius of 0.5 generates a hollow cylinder that decreases the solid cylinder volume with 25% and generates a *SM* of 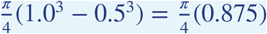. This is a reduction of approximately 13% of *SM*. Many theropods, however, have hollow cores within long bones that comprise 50% or more of the volume of the long bone. For a hollow core equal to 50% of outer cylinder volume, the inner radius is 0.707. The section modulus, calculated by subtracting *SM* for a cylinder of radius 0.707 from the full cylinder of radius 1.0, is 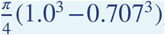, or 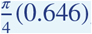, a reduction of approximately 35% of the *SM* of a solid cylinder.

Thus, infilling of hind limb bones in *S. aegyptiacus* may have increased bending strength by as much as 35%. The femur in *S. aegyptiacus*, in particular, may have been subjected to substantial bending forces. The attachment flange for the caudofemoralis muscle is hypertrophied, occupying almost one-third of the length of the femoral shaft.

## Appendix 2

### Extant and extinct comparative materials

We used references in the literature, an extant amphibian and reptile specimens, and a mosasaur to estimate muscle mass in the flesh reconstruction of *S. aegyptiacus* and to measure caudal centrum proportions along the tail (***Figure 2***, ***Figure 4****D*; Appendix 2 ***Table 1***, ***Table 2***).

**Appendix 2 Table 1.** Extant newt and squamate specimens used for muscle mass estimation and for logging centrum proportions along the tail. Specimen number hyperlinks go to Morphosource specimen pages with CT scans and 3D models.

| Specimen No. | Taxon | Sex | Type | Purpose |
| --- | --- | --- | --- | --- |
| FMNH 84926 | *Triturus cristatus* | Male | Wet | Caudal centrum ratios, muscle mass estimation |
| FMNH 57512 | *Intellagama lesueurii* | Male | Wet | Muscle mass estimation |
| FMNH 22389 | *Intellagama lesueurii* | Male | Skeleton | Caudal centrum ratios, muscle mass estimation |
| KU 314941 | *Hydrosaurus amboinensis* | Unknown; low neural spines (female/immature male) | Wet | Caudal centrum ratios |
| FMNH 52698 | *Hydrosaurus pustulatus* | Male | Wet | Muscle mass estimation |
| FMNH 236131 | *Hydrosaurus pustulatus* | Unknown; low neural spines (female/immature male) | Skeleton | Muscle mass estimation |
| FMNH 112993 | *Basiliscus plumifrons* | Male | Wet | Muscle mass estimation |
| FMNH 257162 | *Basiliscus plumifrons* | Male | Skeleton | Additional high resolution visualization |
| UMMZ 121461 | *Basiliscus basiliscus* | Male | Wet | Caudal centrum ratios |
| UF 41558 | *Amblyrhynchus cristatus* | Unknown | Wet | Caudal centrum ratios |
| FMNH 22042 | *Amblyrhynchus cristatus* | Unknown; ?male (tallest caudal neural spines in FMNH collection) | Skeleton | Muscle mass estimation |
| UF 21461 | *Alligator mississippiensis* | Unknown; juvenile | Wet | X-ray image of caudal centra in tail; caudal centrum ratios |

**Appendix 2 Table 2.**
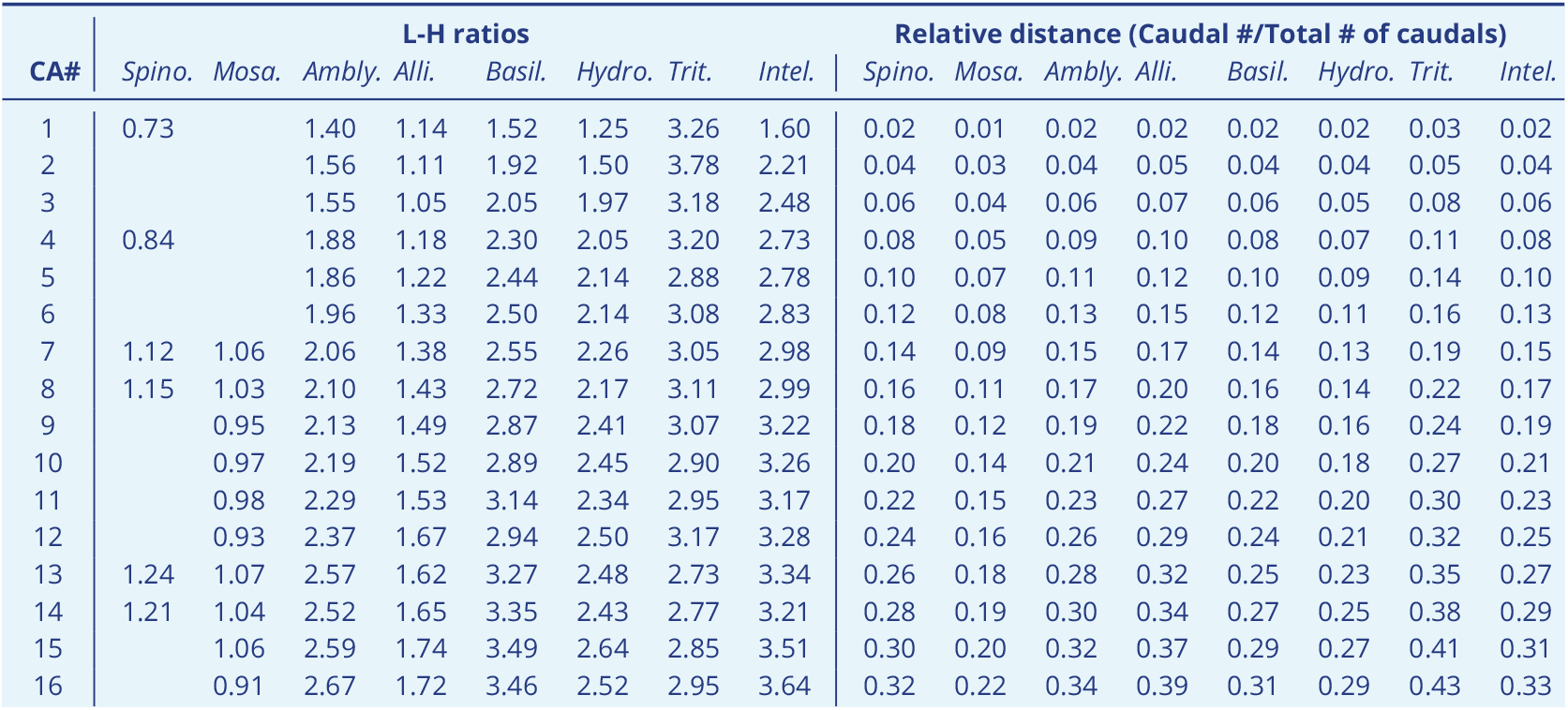
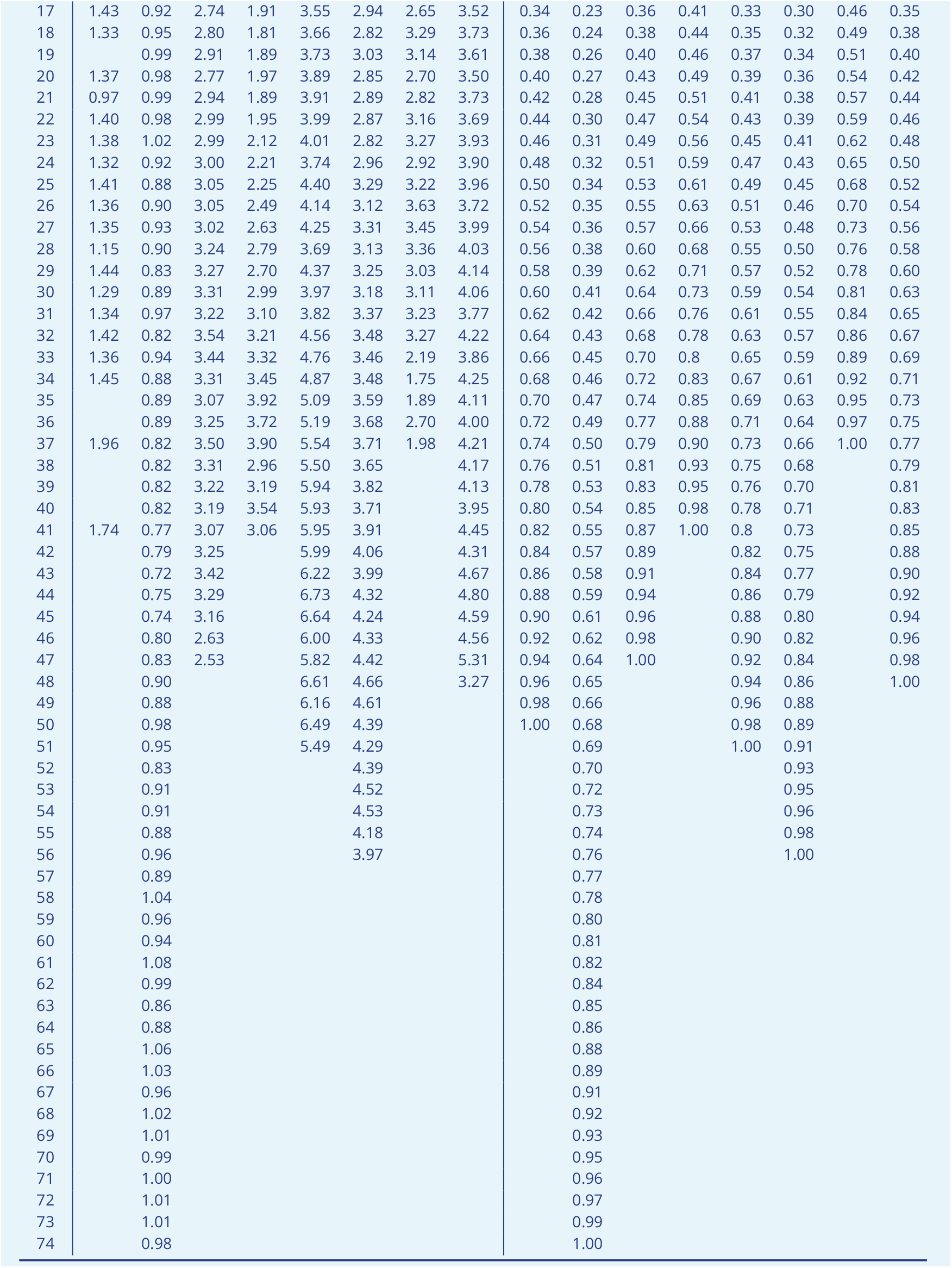
Caudal centrum ratios along the tail in *S. aegyptiacus, Mosasaurus*, and extant semiaquatic amphibians and reptiles (***Figure 4****D*).

| CA# | L-H ratios |  |  |  |  |  |  |  | Relative distance (Caudal #/Total # of caudals) |  |  |  |  |  |  |  |
| --- | --- | --- | --- | --- | --- | --- | --- | --- | --- | --- | --- | --- | --- | --- | --- | --- |
|  | Spino. | Mosa. | Ambly. | Alli. | Basil. | Hydro. | Trit. | Intel. | Spino. | Mosa. | Ambly. | Alli. | Basil. | Hydro. | Trit. | Intel. |
| 1 | 0.73 |  | 1.40 | 1.14 | 1.52 | 1.25 | 3.26 | 1.60 | 0.02 | 0.01 | 0.02 | 0.02 | 0.02 | 0.02 | 0.03 | 0.02 |
| 2 |  |  | 1.56 | 1.11 | 1.92 | 1.50 | 3.78 | 2.21 | 0.04 | 0.03 | 0.04 | 0.05 | 0.04 | 0.04 | 0.05 | 0.04 |
| 3 |  |  | 1.55 | 1.05 | 2.05 | 1.97 | 3.18 | 2.48 | 0.06 | 0.04 | 0.06 | 0.07 | 0.06 | 0.05 | 0.08 | 0.06 |
| 4 | 0.84 |  | 1.88 | 1.18 | 2.30 | 2.05 | 3.20 | 2.73 | 0.08 | 0.05 | 0.09 | 0.10 | 0.08 | 0.07 | 0.11 | 0.08 |
| 5 |  |  | 1.86 | 1.22 | 2.44 | 2.14 | 2.88 | 2.78 | 0.10 | 0.07 | 0.11 | 0.12 | 0.10 | 0.09 | 0.14 | 0.10 |
| 6 |  |  | 1.96 | 1.33 | 2.50 | 2.14 | 3.08 | 2.83 | 0.12 | 0.08 | 0.13 | 0.15 | 0.12 | 0.11 | 0.16 | 0.13 |
| 7 | 1.12 | 1.06 | 2.06 | 1.38 | 2.55 | 2.26 | 3.05 | 2.98 | 0.14 | 0.09 | 0.15 | 0.17 | 0.14 | 0.13 | 0.19 | 0.15 |
| 8 | 1.15 | 1.03 | 2.10 | 1.43 | 2.72 | 2.17 | 3.11 | 2.99 | 0.16 | 0.11 | 0.17 | 0.20 | 0.16 | 0.14 | 0.22 | 0.17 |
| 9 |  | 0.95 | 2.13 | 1.49 | 2.87 | 2.41 | 3.07 | 3.22 | 0.18 | 0.12 | 0.19 | 0.22 | 0.18 | 0.16 | 0.24 | 0.19 |
| 10 |  | 0.97 | 2.19 | 1.52 | 2.89 | 2.45 | 2.90 | 3.26 | 0.20 | 0.14 | 0.21 | 0.24 | 0.20 | 0.18 | 0.27 | 0.21 |
| 11 |  | 0.98 | 2.29 | 1.53 | 3.14 | 2.34 | 2.95 | 3.17 | 0.22 | 0.15 | 0.23 | 0.27 | 0.22 | 0.20 | 0.30 | 0.23 |
| 12 |  | 0.93 | 2.37 | 1.67 | 2.94 | 2.50 | 3.17 | 3.28 | 0.24 | 0.16 | 0.26 | 0.29 | 0.24 | 0.21 | 0.32 | 0.25 |
| 13 | 1.24 | 1.07 | 2.57 | 1.62 | 3.27 | 2.48 | 2.73 | 3.34 | 0.26 | 0.18 | 0.28 | 0.32 | 0.25 | 0.23 | 0.35 | 0.27 |
| 14 | 1.21 | 1.04 | 2.52 | 1.65 | 3.35 | 2.43 | 2.77 | 3.21 | 0.28 | 0.19 | 0.30 | 0.34 | 0.27 | 0.25 | 0.38 | 0.29 |
| 15 |  | 1.06 | 2.59 | 1.74 | 3.49 | 2.64 | 2.85 | 3.51 | 0.30 | 0.20 | 0.32 | 0.37 | 0.29 | 0.27 | 0.41 | 0.31 |
| 16 |  | 0.91 | 2.67 | 1.72 | 3.46 | 2.52 | 2.95 | 3.64 | 0.32 | 0.22 | 0.34 | 0.39 | 0.31 | 0.29 | 0.43 | 0.33 |
| 17 | 1.43 | 0.92 | 2.74 | 1.91 | 3.55 | 2.94 | 2.65 | 3.52 | 0.34 | 0.23 | 0.36 | 0.41 | 0.33 | 0.30 | 0.46 | 0.35 |
| 18 | 1.33 | 0.95 | 2.80 | 1.81 | 3.66 | 2.82 | 3.29 | 3.73 | 0.36 | 0.24 | 0.38 | 0.44 | 0.35 | 0.32 | 0.49 | 0.38 |
| 19 |  | 0.99 | 2.91 | 1.89 | 3.73 | 3.03 | 3.14 | 3.61 | 0.38 | 0.26 | 0.40 | 0.46 | 0.37 | 0.34 | 0.51 | 0.40 |
| 20 | 1.37 | 0.98 | 2.77 | 1.97 | 3.89 | 2.85 | 2.70 | 3.50 | 0.40 | 0.27 | 0.43 | 0.49 | 0.39 | 0.36 | 0.54 | 0.42 |
| 21 | 0.97 | 0.99 | 2.94 | 1.89 | 3.91 | 2.89 | 2.82 | 3.73 | 0.42 | 0.28 | 0.45 | 0.51 | 0.41 | 0.38 | 0.57 | 0.44 |
| 22 | 1.40 | 0.98 | 2.99 | 1.95 | 3.99 | 2.87 | 3.16 | 3.69 | 0.44 | 0.30 | 0.47 | 0.54 | 0.43 | 0.39 | 0.59 | 0.46 |
| 23 | 1.38 | 1.02 | 2.99 | 2.12 | 4.01 | 2.82 | 3.27 | 3.93 | 0.46 | 0.31 | 0.49 | 0.56 | 0.45 | 0.41 | 0.62 | 0.48 |
| 24 | 1.32 | 0.92 | 3.00 | 2.21 | 3.74 | 2.96 | 2.92 | 3.90 | 0.48 | 0.32 | 0.51 | 0.59 | 0.47 | 0.43 | 0.65 | 0.50 |
| 25 | 1.41 | 0.88 | 3.05 | 2.25 | 4.40 | 3.29 | 3.22 | 3.96 | 0.50 | 0.34 | 0.53 | 0.61 | 0.49 | 0.45 | 0.68 | 0.52 |
| 26 | 1.36 | 0.90 | 3.05 | 2.49 | 4.14 | 3.12 | 3.63 | 3.72 | 0.52 | 0.35 | 0.55 | 0.63 | 0.51 | 0.46 | 0.70 | 0.54 |
| 27 | 1.35 | 0.93 | 3.02 | 2.63 | 4.25 | 3.31 | 3.45 | 3.99 | 0.54 | 0.36 | 0.57 | 0.66 | 0.53 | 0.48 | 0.73 | 0.56 |
| 28 | 1.15 | 0.90 | 3.24 | 2.79 | 3.69 | 3.13 | 3.36 | 4.03 | 0.56 | 0.38 | 0.60 | 0.68 | 0.55 | 0.50 | 0.76 | 0.58 |
| 29 | 1.44 | 0.83 | 3.27 | 2.70 | 4.37 | 3.25 | 3.03 | 4.14 | 0.58 | 0.39 | 0.62 | 0.71 | 0.57 | 0.52 | 0.78 | 0.60 |
| 30 | 1.29 | 0.89 | 3.31 | 2.99 | 3.97 | 3.18 | 3.11 | 4.06 | 0.60 | 0.41 | 0.64 | 0.73 | 0.59 | 0.54 | 0.81 | 0.63 |
| 31 | 1.34 | 0.97 | 3.22 | 3.10 | 3.82 | 3.37 | 3.23 | 3.77 | 0.62 | 0.42 | 0.66 | 0.76 | 0.61 | 0.55 | 0.84 | 0.65 |
| 32 | 1.42 | 0.82 | 3.54 | 3.21 | 4.56 | 3.48 | 3.27 | 4.22 | 0.64 | 0.43 | 0.68 | 0.78 | 0.63 | 0.57 | 0.86 | 0.67 |
| 33 | 1.36 | 0.94 | 3.44 | 3.32 | 4.76 | 3.46 | 2.19 | 3.86 | 0.66 | 0.45 | 0.70 | 0.8 | 0.65 | 0.59 | 0.89 | 0.69 |
| 34 | 1.45 | 0.88 | 3.31 | 3.45 | 4.87 | 3.48 | 1.75 | 4.25 | 0.68 | 0.46 | 0.72 | 0.83 | 0.67 | 0.61 | 0.92 | 0.71 |
| 35 |  | 0.89 | 3.07 | 3.92 | 5.09 | 3.59 | 1.89 | 4.11 | 0.70 | 0.47 | 0.74 | 0.85 | 0.69 | 0.63 | 0.95 | 0.73 |
| 36 |  | 0.89 | 3.25 | 3.72 | 5.19 | 3.68 | 2.70 | 4.00 | 0.72 | 0.49 | 0.77 | 0.88 | 0.71 | 0.64 | 0.97 | 0.75 |
| 37 | 1.96 | 0.82 | 3.50 | 3.90 | 5.54 | 3.71 | 1.98 | 4.21 | 0.74 | 0.50 | 0.79 | 0.90 | 0.73 | 0.66 | 1.00 | 0.77 |
| 38 |  | 0.82 | 3.31 | 2.96 | 5.50 | 3.65 |  | 4.17 | 0.76 | 0.51 | 0.81 | 0.93 | 0.75 | 0.68 |  | 0.79 |
| 39 |  | 0.82 | 3.22 | 3.19 | 5.94 | 3.82 |  | 4.13 | 0.78 | 0.53 | 0.83 | 0.95 | 0.76 | 0.70 |  | 0.81 |
| 40 |  | 0.82 | 3.19 | 3.54 | 5.93 | 3.71 |  | 3.95 | 0.80 | 0.54 | 0.85 | 0.98 | 0.78 | 0.71 |  | 0.83 |
| 41 | 1.74 | 0.77 | 3.07 | 3.06 | 5.95 | 3.91 |  | 4.45 | 0.82 | 0.55 | 0.87 | 1.00 | 0.8 | 0.73 |  | 0.85 |
| 42 |  | 0.79 | 3.25 |  | 5.99 | 4.06 |  | 4.31 | 0.84 | 0.57 | 0.89 |  | 0.82 | 0.75 |  | 0.88 |
| 43 |  | 0.72 | 3.42 |  | 6.22 | 3.99 |  | 4.67 | 0.86 | 0.58 | 0.91 |  | 0.84 | 0.77 |  | 0.90 |
| 44 |  | 0.75 | 3.29 |  | 6.73 | 4.32 |  | 4.80 | 0.88 | 0.59 | 0.94 |  | 0.86 | 0.79 |  | 0.92 |
| 45 |  | 0.74 | 3.16 |  | 6.64 | 4.24 |  | 4.59 | 0.90 | 0.61 | 0.96 |  | 0.88 | 0.80 |  | 0.94 |
| 46 |  | 0.80 | 2.63 |  | 6.00 | 4.33 |  | 4.56 | 0.92 | 0.62 | 0.98 |  | 0.90 | 0.82 |  | 0.96 |
| 47 |  | 0.83 | 2.53 |  | 5.82 | 4.42 |  | 5.31 | 0.94 | 0.64 | 1.00 |  | 0.92 | 0.84 |  | 0.98 |
| 48 |  | 0.90 |  |  | 6.61 | 4.66 |  | 3.27 | 0.96 | 0.65 |  |  | 0.94 | 0.86 |  | 1.00 |
| 49 |  | 0.88 |  |  | 6.16 | 4.61 |  |  | 0.98 | 0.66 |  |  | 0.96 | 0.88 |  |  |
| 50 |  | 0.98 |  |  | 6.49 | 4.39 |  |  | 1.00 | 0.68 |  |  | 0.98 | 0.89 |  |  |
| 51 |  | 0.95 |  |  | 5.49 | 4.29 |  |  |  | 0.69 |  |  | 1.00 | 0.91 |  |  |
| 52 |  | 0.83 |  |  |  | 4.39 |  |  |  | 0.70 |  |  |  | 0.93 |  |  |
| 53 |  | 0.91 |  |  |  | 4.52 |  |  |  | 0.72 |  |  |  | 0.95 |  |  |
| 54 |  | 0.91 |  |  |  | 4.53 |  |  |  | 0.73 |  |  |  | 0.96 |  |  |
| 55 |  | 0.88 |  |  |  | 4.18 |  |  |  | 0.74 |  |  |  | 0.98 |  |  |
| 56 |  | 0.96 |  |  |  | 3.97 |  |  |  | 0.76 |  |  |  | 1.00 |  |  |
| 57 |  | 0.89 |  |  |  |  |  |  |  | 0.77 |  |  |  |  |  |  |
| 58 |  | 1.04 |  |  |  |  |  |  |  | 0.78 |  |  |  |  |  |  |
| 59 |  | 0.96 |  |  |  |  |  |  |  | 0.80 |  |  |  |  |  |  |
| 60 |  | 0.94 |  |  |  |  |  |  |  | 0.81 |  |  |  |  |  |  |
| 61 |  | 1.08 |  |  |  |  |  |  |  | 0.82 |  |  |  |  |  |  |
| 62 |  | 0.99 |  |  |  |  |  |  |  | 0.84 |  |  |  |  |  |  |
| 63 |  | 0.86 |  |  |  |  |  |  |  | 0.85 |  |  |  |  |  |  |
| 64 |  | 0.88 |  |  |  |  |  |  |  | 0.86 |  |  |  |  |  |  |
| 65 |  | 1.06 |  |  |  |  |  |  |  | 0.88 |  |  |  |  |  |  |
| 66 |  | 1.03 |  |  |  |  |  |  |  | 0.89 |  |  |  |  |  |  |
| 67 |  | 0.96 |  |  |  |  |  |  |  | 0.91 |  |  |  |  |  |  |
| 68 |  | 1.02 |  |  |  |  |  |  |  | 0.92 |  |  |  |  |  |  |
| 69 |  | 1.01 |  |  |  |  |  |  |  | 0.93 |  |  |  |  |  |  |
| 70 |  | 0.99 |  |  |  |  |  |  |  | 0.95 |  |  |  |  |  |  |
| 71 |  | 1.00 |  |  |  |  |  |  |  | 0.96 |  |  |  |  |  |  |
| 72 |  | 1.01 |  |  |  |  |  |  |  | 0.97 |  |  |  |  |  |  |
| 73 |  | 1.01 |  |  |  |  |  |  |  | 0.99 |  |  |  |  |  |  |
| 74 |  | 0.98 |  |  |  |  |  |  |  | 1.00 |  |  |  |  |  |  |

## Appendix 3

### Crocodylian foot paddle size and scaling

We measured forefoot, hind foot, and tail areas in five species of extant crocodylians by photographing appendages of individuals in captivity (Appendix 3 ***Table 1***).

**Appendix 3 Table 1.**
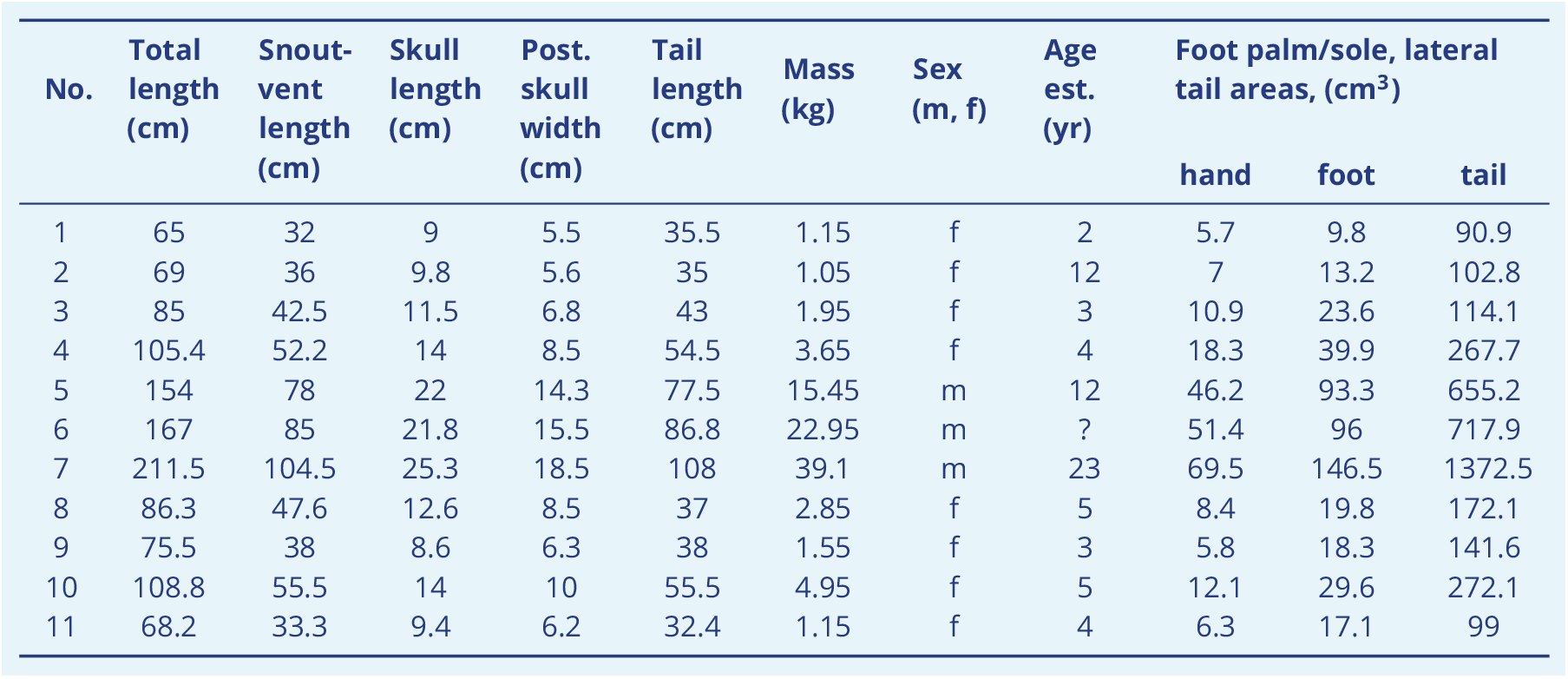
Forefoot, hind foot and tail area and other data in five species of extant crocodylians. American alligator (1-7, *Alligator mississippiensis*); Schneider’s dwarf crocodile (8, *Paleosuchus trigonatus*); Broad-snouted caiman (9, *Caiman latirostris*); Spectacled caiman (10, *Caiman crocodilus*); African dwarf crocodile (11, *Osteolamus tetraspis*).

## Appendix 4

### Inland fossils referable to *Spinosaurus*

Inland fossils referable to *Spinosaurus* sp. were discovered in 1970 at the locality Gara Samani in the Béchar Basin of Algeria (Appendix 4 ***Table 1***). The most complete specimen is a snout (MNHN SAM 124) comparable in size to subadult *S. aegyptiacus* and eventually described as *S. maroccanus* (***Taquet and Russell, 1998***). We consider its specific status in doubt. *S. maroccanus* is based on isolated vertebrae from the Kem Kem Group in Morocco (***Russell, 1996***) and has recently has been reduced to a junior synonym of *S. aegyptiacus* (***Ibrahim et al., 2020b***).

**Appendix 4 Table 1.**
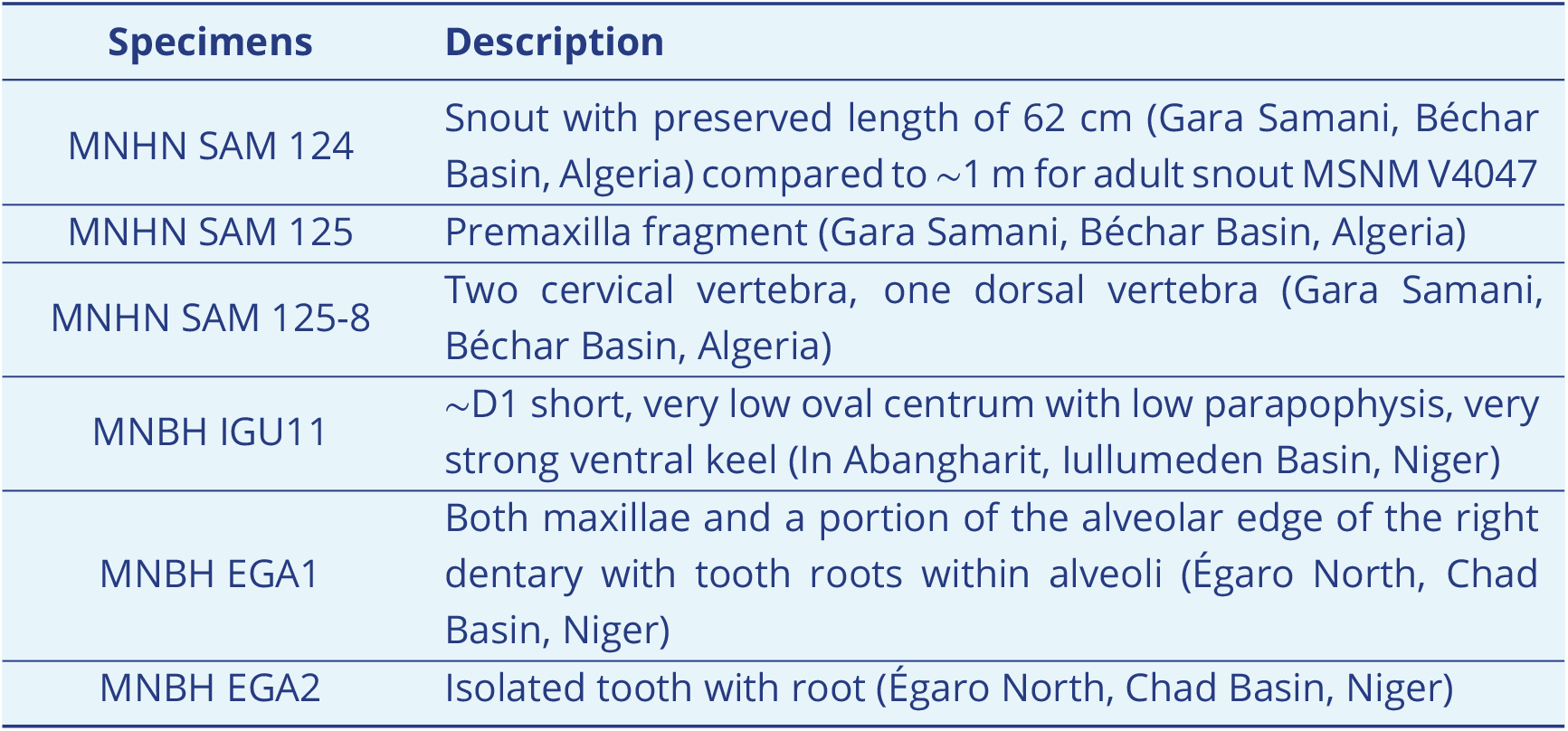
Fossil material referable to *Spinosaurus* from inland basins in Algeria and Niger.

Inland fossils referable to *Spinosaurus* sp. come from two areas of outcrop of the Cenomanianage Echkar Formation in Niger (Appendix 4 ***Table 1***). The first locality is near In Abangharit northeast of Agadez that yielded isolated teeth and an anterior dorsal centrum initially referred to *Carcharodontosaurus iguidensis* (Appendix 4 ***Figure 1***). Reassigned to *Spinosaurus* sp., this vertebra is very close in form to the first dorsal centrum of *S. aegyptiacus* from Morocco (***Ibrahim et al., 2020b***, Fig. 128A-D) and Egypt (***Stromer, 1934***, Pl. 1, Fig. 2), although differences suggest it may pertain to a distinct species. For the time being, reference is made only to the genus *Spinosaurus*.

**Appendix 4 Figure 1.**
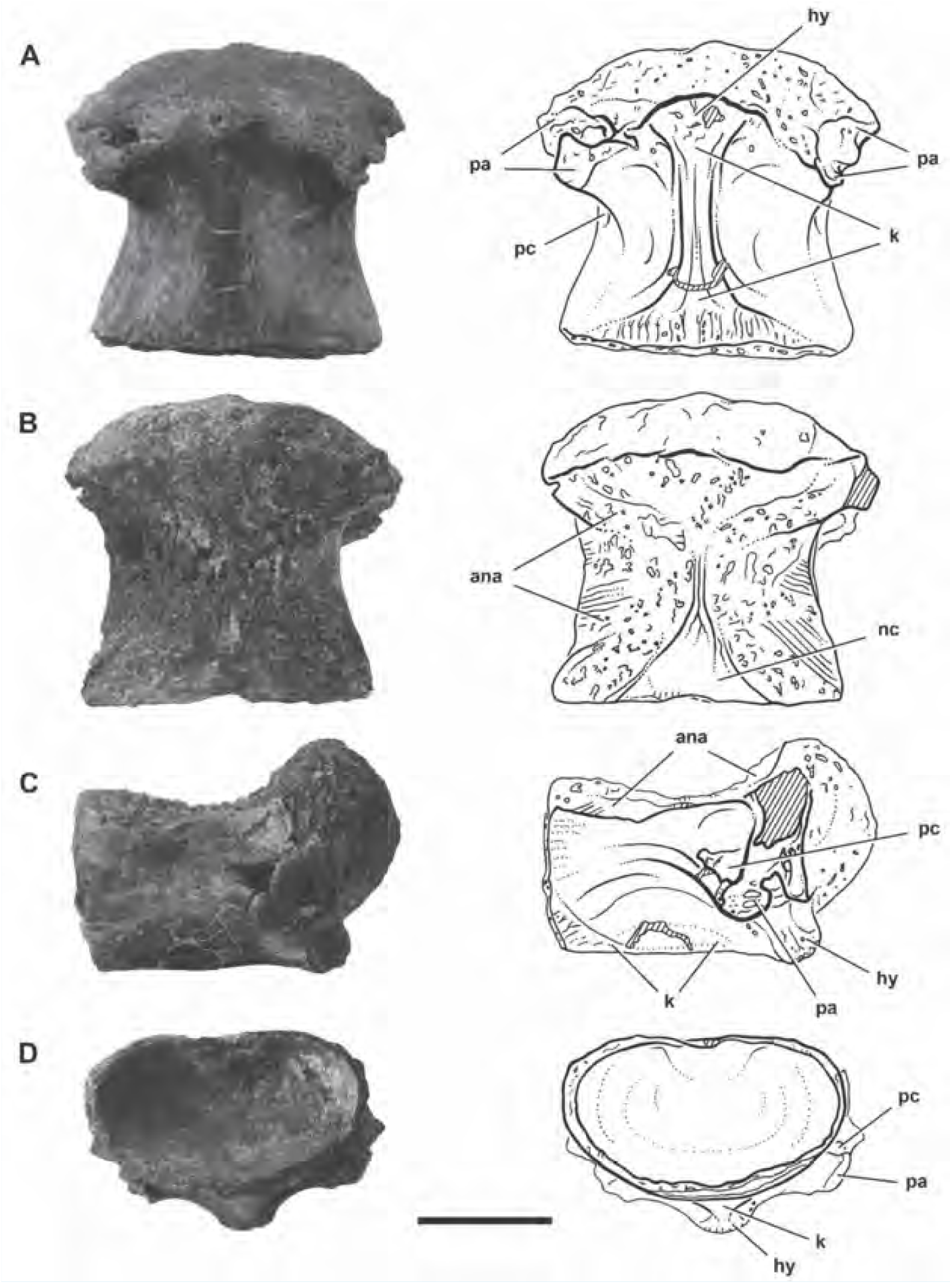
Anterior dorsal centrum referable to *Spinosaurus* sp. (from (***Brusatte and Sereno, 2007***, Figure 9)). Photographs and line drawings of an anterior dorsal centrum (MNBH IGU11) from the Echkar Formation (Cenomanian) of Niger in ventral **(A)** dorsal **(B)**, right lateral **(C)** and posterior **(D)** views. Cross-hatching indicates broken bone. ana, articular surface for the neural arch; hy, hypapophysis; k, ventral keel; nc, neural canal; pa, parapophysis; pc, pleurocoel. Scale bar, 5 cm.

In 2019 a fragmentary snout preserving most of the right and left maxillae of *Spinosaurus* sp. was recovered from a second locality southeast of Agadez also in the Echkar Formation (***Figure 7****B*). The material also includes a partial right dentary (MNBH EGA1) with subconical crowns and long tapering roots (MNBH EGA2). The jaw bones, which closely resemble *S. aegyptiacus* although possibly a new species, are comparable in size to the subadult holotypic and neotypic skeletons, or about 75% of the size of the large snout recovered in Morocco (***Dal Sasso et al., 2005***). They were found in overbank deposits near the remains of rebbachisaurid and titanosaurian sauropods and evidence of a vertebrate fauna common to Cenomanian sites across northern Africa (including *Carcharodontosaurus*, lungfish tooth plates, sawfish rostral teeth, etc.).

## Appendix 5

### Phylogenetic analysis of Spinosauroidea

#### Phylogenetic analysis

A phylogenetic analysis of Spinosauroidea was performed to examine phylogenetic patterns within the clade and survey the distribution and evolution of putative display and semiaquatic features. The analysis included 14 terminal taxa (8 spinosaurids) scored for 120 characters with *Allosaurus fragilis* and *Ceratosaurus nasicornis* as successive outgroups and was analyzed with TNT set for a heuristic search (1000 Wagner tree replicates, 10 TBR; Goloboff et al., 2008). The analysis yielded a single most-parsimonious tree of 156 steps (Appendix 5 ***Figure 1***).

**Appendix 5 Figure 1.**
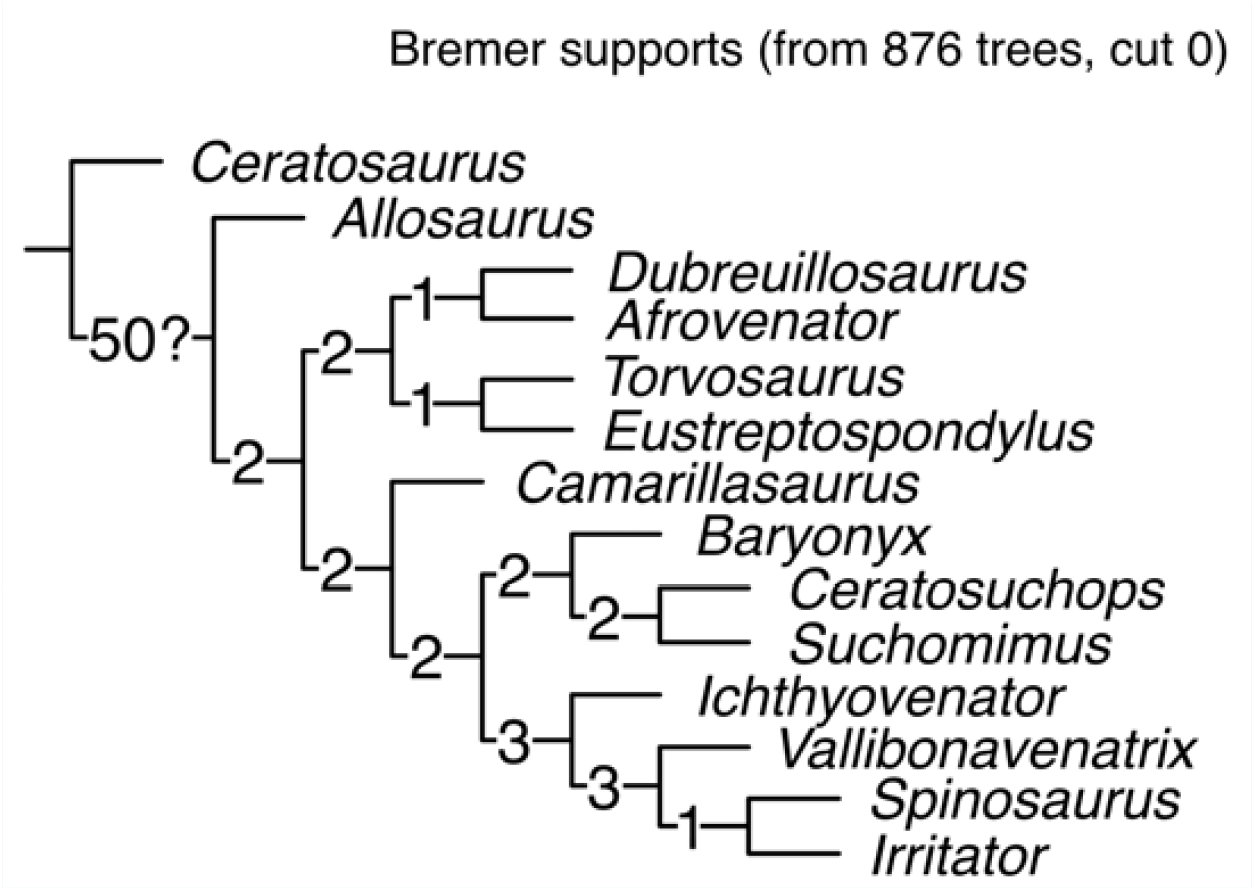
Phylogenetic tree for Spinosauroidea. Single most-parsimonious tree for 12 spinosauroid terminal taxa (8 spinosaurids) and 120 characters split evenly between the cranium (49%) and postcranium (51%), showing decay values (CI = 0.81, RI = 0.86).

Although ***Sales and Schultz (2017***) were unable to resolve a basal polytomy involving *Baryonyx, Suchomimus* and spinosaurines (*Irritator, Spinosaurus*), other analyses follow ***Sereno et al. (1998***) splitting spinosaurid theropods into the subclades Baryonychinae and Spinosaurinae (***Allain et al., 2012***; ***Carrano et al., 2012***; ***Arden et al., 2019***; ***Malafaia et al., 2018***; ***Barker et al., 2021***). We obtain this result again here in an updated analysis of Spinosauroidea based on a dataset of 120 characters, 24 of which are newly introduced. Larger-scale analyses that incorporate theropods far afield and non-applicable character evidence do not appear to have added any clarity or insights to an understanding of ingroup relationships and morphological character change within Spinosauroidea.

*Camarillasaurus cirugedae* was originally described as a ceratosaur (***Sánchez-Hernández and Benton, 2012***) and later reinterpreted as either a megalosaurid or spinosaurid (***Rauhut et al., 2019***; ***Malafaia et al., 2018***; ***Samathi et al., 2021***). Our analysis tentatively resolves *Camarillasaurus* as the most basal spinosaurid, although further study and additional remains of this fragmentary taxon are needed.

Fossil material from the Wessex Formation on the Isle of Wight was described recently as two new baryonychines, *Ceratosuchops inferodios* and *Riperovenator milnerae* (***Barker et al., 2021***), which differ in minor details where they overlap. Both are close in form to *Suchomimus tenerensis*. Only the premaxillae and portions of the braincase of the holotypic specimens of the new taxa overlap, and some doubt exists regarding their association as single specimens, as neither hypodigm were found in association from single sites. The distinguishing features in the premaxillae (low narial tuberosity in one but not preserved in the other) and in the shape/depth of fossae, relative thickness of laminae and other minutiae of the braincase could well be due to individual variation (***Chure and Madsen, 1996***), as several of these features seem to occur on one or the other side of a well-preserved braincase of *Suchomimus tenerensis* (MNBH GAD43) or are impossible to evaluate effectively. The configuration of the orbital margin and form of the swollen postorbital brow is similar in *Suchomimus tenerensis* and *Ceratosuchops*, which was cited as distinguishing feature of the latter. We tentatively score as a single taxon the Wessex spinosaurid material (shown as *Ceratosuchops* on the phylogeny), which is resolved as the sister taxon to *Suchomimus tenerensis* (Appendix 5 ***Figure 2***).

**Appendix 5 Figure 2.**
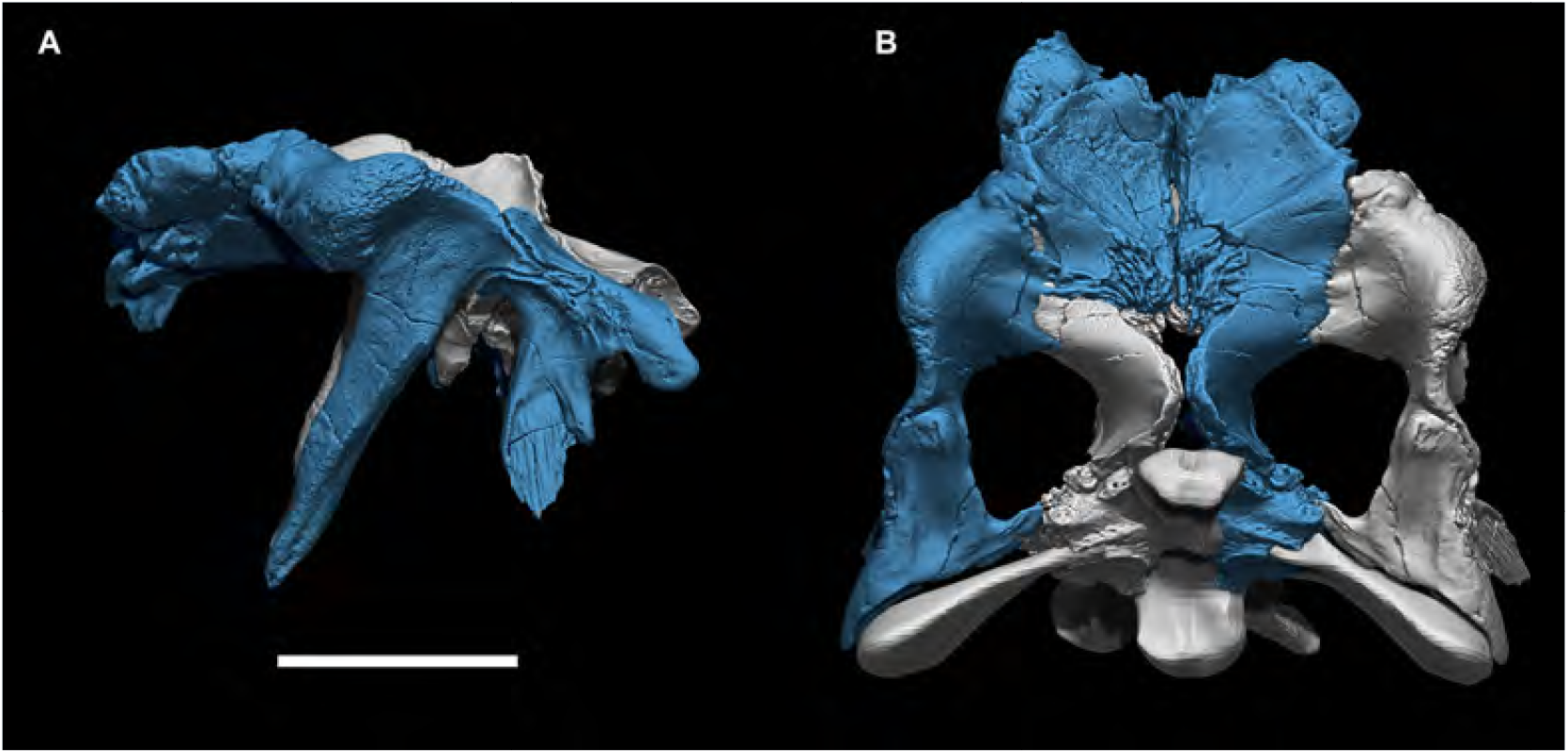
Posterior skull roof of the baryonychine spinosaurid *Suchomimus tenerensis*. Composite restoration of the posterior skull roof of *Suchomimus tenerensis* in **(A)** lateral and **(B)** dorsal views showing a swollen postorbital brow and narrow orbital notch limiting the frontal orbital margin. Scale = 10 cm.

The analysis positions *Ichthyovenator* (***Allain et al., 2012***) and *Vallibonavenatrix* (***Malafaia et al., 2018***) as successively closer to the spinosaurines *Irritator* and *Spinosaurus*. Postcranial remains from the Romualdo Formation of Brazil (***Aureliano et al., 2018***), which may pertain to the Brazilian genus *Irritator*, show derived spinosaurine features (e.g., posterior dorsal neural spines narrower than centrum length, iliac blade proportionately low, pubic process of the ilium ventrally directed). *Irritator* and *Spinosaurus* share straight tooth crowns, spaced tooth positions, more posteriorly positioned external nares, and an oval quadrate head.

##### Character list

**Skull**

1. Snout (preorbital region of skull), length relative to antorbital fenestra length: less (0) or more (1) than three times. (***Sereno et al., 1998***, char. 7)
2. Jaws (premaxillae and dentary), anterior end, form: convergent (0); expanded into a premaxillary/ dentary rosette (1). (***Sereno et al., 1998***, char. 6)
3. Premaxilla, form of premaxilla-nasal suture: V-shaped (0); W-shaped (1). (***Carrano et al., 2012***, char. 5)
4. Premaxilla, external nares, proportions and position: shorter than premaxilla ventral to nares, angle between anterior and alveolar margins > 75° (0); longer than body ventral to nares (1). (***Carrano et al., 2012***, char. 6)
5. Premaxillae, inter-premaxillary suture at maturity, form: open (0); fused (1). (***Sereno et al., 1998***, char. 10)
6. Premaxilla-maxilla articulation, form: scarf or butt joint (0); interlocking (1). (***Sereno et al., 1998***, char. 11)
7. Premaxilla-maxilla suture, lateral surface, subnarial foramen, shape: foramen (0); dorsoventrally directed channel (1). (***Carrano et al., 2012***, char. 10)
8. Premaxilla, ventral margin, shape in lateral view: straight to convex (0); concave (1). (***Cau et al., 2017***, char. 1485)
9. Premaxilla, lateral and dorsal surface, extensive pitting of neurovascular foramina: absent (0); present (1). (***Barker et al., 2021***, char. 1796)
10. Maxilla, anterior ramus, length relative to maximum depth: 70% (0); 100% or more (1). (***Sereno et al., 1998***, char. 1)
11. Maxilla, contribution to the narial fossa: no contribution (0); contributes partially or totally (1). (modified from ***Longrich and Currie (2009***), char 7).
12. Maxilla, antorbital fossa, width of ventral margin relative to depth of posterior ramus of maxilla: more (0) or less (1) than 30%. (***Sereno et al., 1998***, char. 40)
13. Maxilla, antorbital fossa, anterior margin: rounded (0); squared (1). (***Carrano et al., 2012***, char. 23)
14. Maxilla, subcircular depression in the anterior corner of the antorbital fossa: absent (0); present (1). (***Sereno et al., 1998***, char. 41)
15. Maxilla, anteromedial process, shape: fluted prong (0); plate (1). (***Sereno et al., 1998***, char. 12)
16. Maxilla, anteromedial process, anterior extension: as far as (0) or far anterior to (1) the anterior margin of the maxilla. (***Sereno et al., 1998***, char. 13)
17. Maxilla, antorbital fossa, size relative to orbit: larger (0); smaller (1). (***Sereno et al., 1998***, char. 9)
18. Maxilla, anteroventral margin, curvature: not curved (0); dorso-medially curved (1). (modified from ***Cau et al. (2017***), char 731; ***Tykoski (2005***))
19. Nasal, posterior process, overlap of frontal in articulation: absent or limited (0); extensive, in particular medially, on almost or all the process dorsal surface (1). (modified from ***Cau et al. (2017***), char. 1500)
20. Jugal, posterior ramus, depth relative to orbital ramus: less (0); more (1). (***Sereno et al., 1998***, char. 43)
21. Lacrimal, anterior and ventral rami, angle of divergence: 75° to 90° (0); 30° to 45° (1). (***Sereno et al., 1998***, char. 15)
22. Lacrimal foramen, position relative to ventral process: near the base (0); at mid-height (1). (***Sereno et al., 1998***, char. 42)
23. Lacrimal, anterior ramus, length relative to ventral ramus: more (0) or less (1) than 65%. (***Sereno et al., 1998***, char. 2)
24. Postorbital, ventral process, cross section of distal half: subcircular (0); U-shaped (1). (***Sereno et al., 1998***, char. 44)
25. Postorbital, supraorbital shelf (boss) formed mostly by palpebral: absent (0); present (1). (modified from ***Carrano et al. (2012***), char. 61)
26. Frontal, postorbital facet, anterior depth: less than 2/5 facet length (0); more than 2/5 facet length (1). (***Barker et al., 2021***, Supplementary 2)
27. Frontal, shape of the lateral margin in dorsal view: describes a smooth transition between the anterior half and the postorbital process (0); an abrupt transition between the anterior half and the postorbital process (1). (***Senter et al., 2010***, char. 44)
28. Prefrontal, boss-like process: absent (0); present (1). (***Barker et al., 2021***, Supplementary 2)
29. Parietal, length: less than 3/4 of the frontal (0); subequal or more than 3/4 of the frontal (1). (***Cau et al., 2017***, char. 78)
30. Quadrate, head, shape: oval (0); subquadrate (1). (***Sereno et al., 1998***, char. 27)
31. Quadrate, foramen, size: foramen (0); broad fenestra (1). (***Sereno et al., 1998***, char. 28)
32. Quadrate, medial foramina adjacent to condyles: absent (0); present (1). (***Carrano et al., 2012***, char. 84)
33. Quadrate, foramen margin, placement: at mid-height or dorsal (0); ventral, close to mandibular condyles (1). (modified from ***Loewen et al. (2013***), char. 158)
34. Basisphenoid, width of the interbasipterygoidal web: thin; 40 percent of occipital condyle width or less (0); thick, more than 40 percent occipital condyle width (1). (modified from ***Barker et al. (2021***), Supplementary 2)
35. Basisphenoid, basipterygoid process, exposure of the ventral surface in lateral view: thick (0); reduced (1). (***Barker et al., 2021***, Supplementary 2)
36. Basisphenoid, basipterygoid process, shape of lateral margin in ventral view: flat or slightly concave (0); convex (1). (***Barker et al., 2021***, Supplementary 2)
37. Basioccipital, position of subcondylar recess: dorsoventrally tall, recess reaches the occipital condyle neck (0); ventrally restricted, surface directly below condyle convex (1). (***Barker et al., 2021***, Supplementary 2)
38. Basioccipital, width of subcondylar recess relative to occipital condyle width: narrow, 0.5 times or less (0); wide, greater than 0.5 times (1). (modified from (***Barker et al., 2021***, Supplementary 2))
39. Basioccipital, thick crests bordering subcondylar recess laterally: present (0); absent (1). (***Barker et al., 2021***, Supplementary 2)
40. Basioccipital, contribution to foramen magnum: large, exoccipitals widely separated (0); reduced, exoccipitals closely placed (1). (***Barker et al., 2021***, Supplementary 2)
41. Basioccipital, proportions relative to basisphenoid (measured along midline ventral to occipital condyle to interbasipterygoidal web), in posterior view: shorter (0); longer (1). (***Barker et al., 2021***, Supplementary 2)
42. Otoccipitals, angle of projection of paroccipital processes: posterolaterally (0); laterally or subhorizontal (1). (***Barker et al., 2021***, Supplementary 2)
43. Splenial foramen, size: small (0); large (1). (***Sereno et al., 1998***, char. 16)
44. Dentary, shape of anterior end in lateral view: blunt and unexpanded (0); dorsoventrally expanded, rounded and slightly upturned (1); squared off in lateral view via anteroventral process (2). (***Carrano et al., 2012***, char. 120)
45. Premaxilla and anterior dentary, interdental septa spacing: regular (0); alternate (alveoli result paired) (1). (modified from ***Cau et al. (2017***), char. 1614) **Dental**
46. Teeth, distal, curvature: present, marked (0); reduced or non-curved (1). (***Sereno et al. (1998***), char. 35; modified after ***Hendrickx et al. (2019***))
47. Teeth, maxillary and dentary, serrations: present (0); absent (1). (***Sereno et al., 1998***, char. 17)
48. Teeth, distal, midcrown cross-section: elliptical (0); circular (1). (***Sereno et al. (1998***), char. 36; modified after ***Hendrickx et al. (2019***))
49. Teeth, distal, crown striations (flutes/apicobasal ridges): absent (0); present (1). (***Sereno et al. (1998***), char. 18; modified after ***Hendrickx et al. (2019***))
50. Teeth, enamel ornamentation: absent (0); present (1). (modified from ***Carrano et al. (2012***), char. 143)
51. Teeth, enamel ornamentation type: extending as bands across labial and lingual tooth surfaces (0); pronounced marginal enamel wrinkles (1); pronounced deeply veined/ anastomous texture (2). (modified from ***Carrano et al. (2012***), char. 143)
52. Teeth, basalmost root shape: broad (0); strongly tapered (1). (***Sereno et al., 1998***, char. 21)
53. Premaxilla, tooth count: 3 to 4 (0); 6 to 7 (1). (***Sereno et al., 1998***, char. 19)
54. Premaxillary tooth 1, size: slightly smaller (0) or much smaller (1) than crowns 2 and 3. (***Sereno et al., 1998***, char. 38)
55. Premaxilla, diastemata within the premaxillary rosette: narrow (0); broad (1). (***Sereno et al., 1998***, char. 39)
56. Maxillary teeth, mid-tooth spacing: adjacent (0); with intervening space/diastemata (1). (***Sereno et al., 1998***, char. 20)
57. Dentary, tooth count: up to 15 (0); ≥ 15 (1). (***Sereno et al., 1998***, char. 26)
58. Dentary teeth, spacing: adjacent (0); with intervening space (1). (***Sereno et al., 1998***, char. 37)
59. Paradental laminae: present (0); absent (1). (***Sereno et al., 1998***, char. 14) **Axial**
60. Cervical vertebrae, middle, shape of anterior pneumatic foramina: round (0); anteroposteriorly elongate (1). (***Carrano et al., 2012***, char. 169)
61. Cervical vertebrae, posteriormost, ventral keel: absent or developed as a weak ridge (0); pronounced, around 1/3 the height of centrum and inset from lateral surfaces (1). NEW
62. Cervical vertebrae, prezygapophyseal facets, elongation related to width: as long as wide (0); longer than wide (1); wider than longer (2). NEW
63. Cervical vertebrae, middle and posterior centra width relative to centra height: taller than wide or round (0); wider than tall (1). NEW
64. Cervical vertebrae, anterior post-axial neural spines in lateral view: longer than tall (0); taller than long (1). (***Cau et al., 2017***, char. 212)
65. Dorsal vertebrae, anterior centra, depth of ventral keel relative to total centrum height: absent or less than ¼ (0); blade-shaped, more than ¼ (1). (***Sereno et al., 1998***, char. 22)
66. Dorsal vertebrae, anterior centra, ventral processes anterior to the keel (hypapophysis): poorly developed (0); strongly developed (1). (Modified by ***Cau et al. (2017***) (char. 225) from ***Rauhut (2003***); ***O’Connor (2009***))
67. Dorsal vertebrae, middle, centra length relative to height: 1.4 < times centrum height (0); ≥ 1.4 times centrum height (1). NEW
68. Dorsal vertebrae, anterior parapophyses, size: less (0) or more (1) than half-depth of anterior facet of centrum. (***Cau et al., 2017***, char. 1740)
69. Dorsal vertebrae, anterior centra, pneumatic foramen, size relative to parapophysis: larger or equal (0); smaller (1). NEW
70. Dorsal vertebrae, anterior, prezygapophyseal facets, elongation related to width: as long as wide (0); longer than wide (1); wider than longer (2). NEW
71. Dorsal vertebrae, mid-posterior, excavated prezygo-para-diapophyseal fossa (prpadf) delimited by the paradiapophyseal lamina (ppdl): absent (0); present (1). NEW (after ***Malafaia et al. (2020***))
72. Dorsal vertebrae, middle and posterior centra, pneumatic foramina: absent (0); present (1). (modified from ***Cau et al. (2017***), char. 1350)
73. Dorsal vertebrae, height of neural spines relative to centrum height: low, < 1.3x (0); moderate, 1.3-1.8x (1); tall, < 1.8x (2). (modified from ***Carrano et al. (2012***), char. 193)
74. Dorsal vertebrae, posterior neural spines, basal webbing: absent (0); present (1). (***Sereno et al., 1998***, char. 24)
75. Dorsal vertebrae, posterior neural spines, accessory centrodiapophyseal lamina: absent (0); present (1). (***Sereno et al., 1998***, char. 25)
76. Dorsal vertebrae, middle and posterior, neural spine antero-posterior length at base relative to centrum length: subequal (0); shorter (1). NEW
77. Dorsal vertebrae, middle-posterior parapophyses, development: distinct and well-developed (0); small, knob-like (1). (***Stromer, 1934***)
78. Dorsal vertebrae, accessory centrodiapophyseal lamina: absent (0); present (1). (modified from ***Benson et al. (2009***), char. 2015)
79. Sacrum, centrum pneumaticity: absent (0); present (1). (modified from ***Carrano et al. (2012***), char. 196)
80. Sacrum, centrum pneumaticity type: pleurocoelous fossae (0); pneumatic foramina (1). (modified from ***Carrano et al. (2012***), char. 196)
81. Sacrum, neural spines: without distal antero-posterior expansion (0); with distal expansion contacting adjacent spines (1). NEW
82. Caudal vertebrae, anterior, morphology of ventral surface: flat (0); groove (1); ridge (2). (***Carrano et al., 2012***, char. 202)
83. Caudal vertebrae, anterior, well-marked spino-diapophyseal lamina: absent (0); present (1). NEW
84. Caudal vertebrae, anterior neural arches, ventral rib laminae: absent (0); present (1). (***Cau et al., 2017***, char. 358)
85. Caudal vertebrae, anterior neural arches, anterolateral surface, deep triangular prezygocostal fossa delimited by two laminae: absent (0); present (1). (***Cau et al. (2017***), char. 1605; modified from ***Brusatte et al. (2010***))
86. Caudal vertebrae, anterior neural arches, hyposphene: absent (0); present (1). (***Cau et al., 2017***, char. 359)
87. Caudal vertebrae, CFL and ilio-ischiocaudalis correlates: transition point at or beyond CA20 (0); transition point around CA15-19 (1). NEW
88. Caudal vertebrae, middle, height of neural spines relative to centrum height: low, ≥ 1.3x (0); moderate, 1.3-2.0x (1); tall, 2.0-4.0x (2); extreme elongation > 4x (3). NEW
89. Caudal vertebrae, distal, height of neural spines relative to centrum height: low, ≥ 1.3x (0); moderate, 1.3-2.0x (1); tall, 2.0-4.0x (2); extreme elongation > 4x (3). NEW
90. Caudal vertebrae, middle, morphology of neural spines: rod-like and posteriorly inclined (0); subrectangular and sheet-like (1); rod-like and vertical (2). (***Carrano et al., 2012***, char. 207)
91. Caudal vertebrae, middle, centrum elongation relative to centrum height: elongated > 1.6x (0); not elongated ≤ 1.6x (1). NEW
92. Caudal vertebrae, middle and posterior, prezygapophyses: elongated, projected beyond the anterior rim of the centrum (0); shortened, barely reaching the anterior rim of the centrum (1). NEW (after observations in ***Samathi et al. (2021***))
93. Chevrons, anterior and posterior longitudinal groove: absent (0); present (1). NEW
94. Chevrons, anterior and posterior surfaces: without distinctive features (0); with longitudinal groove widened as a fossa (1). NEW
95. Chevrons, posterior elongation compared with anterior and middle ones: shorter (0); as elongated (1). NEW
96. Chevrons, anterior process: absent (0); present (1). (***Carrano et al., 2012***, char. 217) **Appendicular**
97. Coracoid, posterior process, shape: low and rounded (0); crescentic (1). (***Sereno et al., 1998***, char. 29)
98. Appendicular bones, marrow cavity: present (0); reduced to barely present (1). NEW
99. Humerus, deltopectoral crest, length relative to humeral length: less (0) or more (1) than 45%. (***Sereno et al., 1998***, char. 3)
100. Humerus, deltopectoral crest, orientation of apex: anterior (0), lateral (1). (***Sereno et al., 1998***, char. 31)
101. Humerus, trochanters, size: low and rounded (0); hypertrophied and pointy (1). (***Sereno et al., 1998***, char. 30)
102. Humerus, internal tuberosity, size: low and rounded (0); hypertrophied (1). (***Sereno et al., 1998***, char. 32)
103. Radius (forearm), length relative to humeral length: more (0) or less (1) than 50%. (***Sereno et al., 1998***, char. 4)
104. Radius, external tuberosity and ulnar internal tuberosity, size: low and rounded (0); hypertrophied (1). (***Sereno et al., 1998***, char. 33)
105. Manual digit I–ungual, length relative to the depth of proximal end: 2.5 (0) or 3 (1) times. (***Sereno et al., 1998***, char. 5)
106. Ilium, ventrolateral development of supraacetabular crest: large/pendant ’hood’ (0); reduced shelf (1). (***Carrano et al., 2012***, char. 267)
107. Ilium, lateral vertical crest dorsal to the acetabulum: absent (0); present (1). (***Rauhut, 2003***, char. 6)
108. Ilium, shape of posterior margin of postacetabular process: convex (0); concave (1); straight (2); with prominent posterodorsal process but lacking posteroventral process (3). (***Carrano et al., 2012***, char. 280)
109. Ilium, orientation of pubic peduncle: mostly ventral (0); mostly anterior or ’kinked’ double facet with anterior and ventral components (1). (***Carrano et al., 2012***, char. 268)
110. Ilium, pubic peduncle length to width ratio: ≤ 1 (0); 1-2 (1); > 2 (2). (modified from ***Carrano et al. (2012***), char. 272)
111. Ilium, brevis fossa, lateral and medial margins, orientation in ventral view and development of fossa: subparallel, narrow fossa (0); posteriorly diverging, expanded fossa (1). (modified by ***Cau et al. (2017***) (char. 592) from ***Holtz (2000***); ***Rauhut (2003***))
112. Puboischiadic plate, foramina/notches: closed along midline, 3 fenestrae (0); open along midline, 1 fenestra (obturator foramen of pubis) and 1-2 notches (1); open along midline, 0 fenestrae, 1-2 notches (2). (***Carrano et al., 2012***, char. 281)
113. Pubis, distal pubic foot, size: moderate to large (0); reduced to a small flange (1). (modified from ***Sereno et al. (1998***), char. 34)
114. Pubis, length relative to ischium: longer (0); subequal or shorter (1). NEW
115. Ischium, distal half, cross section: laminar, strongly mediolaterally compressed (0); robust, rod-like (1). (***Cau et al., 2017***, char. 425)
116. Femur, fourth trochanter position of distalmost end: proximal most 1/3 (0); almost at the half of the femur (1). NEW
117. Femur, oblique ligament groove on posterior surface of head: shallow, groove bounding lip does not extend past posterior surface of head (0); deep, bound medially by well-developed posterior lip (1). (***Carrano et al., 2012***, char. 304)
118. Tibia and/or femur, length compared to posterior dorsal centra length: more (0) or less (1) than five times. (***Cau et al., 2017***, char. 245)
119. Tibia, proximal diaphysis, length-width ratio: smaller than 2 (0); greater than 2 (1). NEW (after ***Samathi et al. (2021***))
120. Pedal ungual phalanges, ventral side: concave (0); flattened (1). NEW

##### Character-taxon matrix

*Ceratosaurus* 0 0 1 0 0 0 0 0 0 0 0 0 0 0 0 0 0 0 0 0 0 0 0 0 0 0 0 0 0 0 0 0 0 0 0 0 0 ? ? 0 0 0 0 0&1 0 0 0 0 0 1 0 0 0 0 ? 0 0 0 0 0 0 ? 0 1 0 0 0 0 ? 0 1 0 0 0 0 0 0 0 0 ? 0 1 0 0 0 1 ? 0 0 2 0 ? 0 ? 0 0 1 0 0 0 0 0 0 0 0 0 0 1 1 0 1 1 0 0 1 0 1 0 0 0

*Allosaurus* 0 0 1 0&1 0 0 0 0 0 0 0 0 1 0 0 0 0 0 0 0 0 0 0 0 0 0 0 0 0 0 0 0 0 0 0 0 0 0 0 0 0 0 0 0&1&2 0 0 0 0 0 0&1&2 0&1&2 0 0 0 ? 0 0 0 0 0&1 0 0&1&2 0 1 0 0 0 0 0 0 0 0 0&1&2 0 0 0 0 0 1 0 0 1 0 0 0 0 0 0&1 0&1 0 0 0 0 ? 0 0 1 0 0 0 1 0 0 0 0 1 0 0 0&1 2 0 2 0 0 1 0 1 0 0 0

*Eustreptospondylus* 0 0 ? 1 0 0 0 0 0 1 0 1 ? 1 0 0 0 0 ? 1 0 1 1 1 0 0 0 0 0 0 0 0 0 ? ? ? 0 0 0 0 ? 0 ? 0 0 0 0 0 0 1 1 0 0 0 ? 0 0 0 0 1 0 ? 0 1 0 0 ? 0 0 0 0 ? 1 0 0 ? ? ? 0 ? 0 1 0 0 0 1 ? ? ? 0 0 ? ? ? ? ? 0 ? 1 0 0 0 ? ? ? 1 ? 3 1 1 ? ? 1 ? ? 0 0&1 ? 0 0

*Torvosaurus* 0 0 ? 1 0 0 0 0 0 1 0 1 0 1 0 0 0 0 ? 1 0 1 1 1 0 ? ? ? ? 0 0 1 0 ? ? ? ? ? ? ? ? ? ? 0 0 0 0 0 0 1 1 0 0 0 ? 0 0 0 0 0 0 0 0 1 0 0 0 0 0 0 0 0&1 1 0 0 0 0 0 1 0 0 1 0 0 1 0 ? 0 ? 0 0 ? 1 0 0 1 0 ? 1 0 0 0 1 0 1 1 0 3 1 1 1 0 1 0 1 0 0 0 0 0

*Afrovenator* 0 ? ? ? ? 0 ? ? ? 1 0 1 1 1 0 0 0 0 ? 1 0 1 0 1 0 ? ? ? ? 0 0 1 ? ? ? ? ? ? ? ? ? ? ? ? ? 0 0 0 0 1 1 ? ? ? ? 0 ? ? ? 1 0 0 0 0 0 0 0 0 0 ? 0 0 0 0 0 0 0 0 ? ? ? ? ? ? ? ? ? 0 0 0 0 0 1 0 0 1 ? ? 0 0 0 0 ? ? 0 1 1 0 1 1 0 ? 1 0 1 0 0 0 ? 0

*Dubreuillosaurus* 0 0 ? 1 0 0 ? 0 0 1 0 1 1 ? 0 0 0 0 ? ? 0 ? 1 ? 0 0 ? ? ? ? ? ? ? ? ? ? 0 0 1 0 ? 0 1 0 0 0 0 0 0 1 1 0 0 0 ? 0 0 0 0 1 0 ? ? ? 0 ? ? ? ? ? ? ? ? ? ? ? ? ? 1 1 ? ? 0 ? ? ? ? ? ? ? 0 0 1 0 0 1 ? ? ? ? ? ? ? ? ? ? ? ? ? ? ? ? ? ? ? ? 0 ? ? ?

*Baryonyx* 1 1 0 2 1 1 1 1 1 1 0 ? ? ? 1 1 1 1 ? ? 1 0 1 ? ? 0 0 ? 1 1 1 0 ? 0 0 0 0 1 0 0 0 1 1 1 0 0 0 1 1 1 2 1 1 0 0 1 1 0 1 1 1 1 0 0 1 0 0 1 0 1 0 0 2 1 1 0 0 1 ? ? ? ? 0 ? ? ? ? ? ? ? ? ? 1 1 ? 0 2 ? 1 1 1 1 1 1 1 1 ? 0 0 1 ? ? 1 ? 0 ? 1 ? ? ?

*Suchomimus* 1 1 0 2 1 1 1 1 1 1 0 ? ? 0 1 1 1 1 1 ? ? ? 1 0 1 1 1 1 1 1 1 0 ? 1 0 0 0 0 1 1 1 0 ? 1 0 0 0 1 1 1 2 1 1 0 0 1 1 0 1 1 1 1 0 0 1 0 0&1 1 0 1 0 0 2 1 1 0 0 1 1 1 1 0 0&1 0 0 0 0 2 ? 0 0 1 1 1 0 0 2 0 1 1 1 1 1 1 1 1 ? 0 1 0 1 0 1 0 0 0 1 0 1 0

*Ichthyovenator* ? ? ? ? ? ? ? ? ? ? ? ? ? ? ? ? ? ? ? ? ? ? ? ? ? ? ? ? ? ? ? ? ? ? ? ? ? ? ? ? ? ? ? ? ? ? ? ? ? ? ? ? ? ? ? ? ? ? ? ? 1 1 1 1 1 1 0 1 1 1 1 0 2 ? 1 1 1 0 ? ? 1 0 0 1 1 0 ? 2 ? 0 0 1 1 1 1 0 ? ? ? ? ? ? ? ? ? 0 ? 2 1 2 ? 1 1 0 ? ? ? ? ? ?

*Camarillasaurus* ? ? ? ? ? ? ? ? ? ? ? ? ? ? ? ? ? ? ? ? ? ? ? ? ? ? ? ? ? ? ? ? ? ? ? ? ? ? ? ? ? ? ? ? ? ? ? 1 0 ? ? ? ? ? ? ? ? ? ? ? ? ? ? ? ? ? ? ? ? ? 0 ? ? ? ? ? ? ? 0 ? ? 1 ? ? ? 1 ? ? ? ? 0 1 1 0 ? 0 ? ? ? ? ? ? ? ? ? ? ? ? ? ? ? ? ? ? ? ? ? ? 1 ?

*Vallibonavenatrix* ? ? ? ? ? ? ? ? ? ? ? ? ? ? ? ? ? ? ? ? ? ? ? ? ? ? ? ? ? ? ? ? ? ? ? ? ? ? ? ? ? ? ? ? ? ? ? ? ? ? ? ? ? ? ? ? ? ? ? 1 1 ? 1 ? 1 ? 1 1 ? ? 1 1 2 ? 1 1 ? 1 1 1 ? 1 1 ? ? 0 ? ? ? ? 1 ? 0 ? ? 0 1 ? ? ? ? ? ? ? ? 1 0 0 1 2 1 ? ? ? 0 ? ? ? ? ?

*Irritator* 1 1 ? 2 1 1 ? ? ? 1 1 ? 0 0 1 1 1 1 1 0 1 ? 1 1 0&1 0 0 1 1 0 1 0 1 0 1 0 0 0 0 0 0 0 ? ? ? 1 1 1 1 1 2 1 ? 1 1 1 0 1 ? ? ? ? ? ? ? ? ? ? ? ? ? 1 2 ? ? 1 ? ? 1 1 1 ? ? ? ? ? ? ? ? 0 1 ? ? ? ? ? ? 1 ? ? ? ? ? ? 1 1 0 0 0 1 1 0&1 ? ? 0 ? ? ? ? ?

*Spinosaurus* 1 1 1 2 1 1 1 1 1 1 1 ? ? ? 1 1 ? 1 1 ? ? ? ? ? ? ? ? ? ? 0 1 0 1 ? ? ? ? ? ? ? ? ? 1 1 1 1 1 1 0&1 1 2 1 1 1 1 1 0 1 1 1 1 1 1 1 1 1 1 1 1 1 1 1 2 0 1 1 1 0 1 1 1 1 1 1 1 0 1 3 3 0 1 1 1 1 1 0 ? 1 ? ? ? ? ? ? 1 1 0 0 0 1 1 1 0 1 0 1 1 1 1 1

*Ceratosuchops* ? 1 0 2 1 1 1 1 1 ? ? ? ? ? ? ? ? ? ? ? ? ? ? 1 1 1 1 1 1 ? ? ? ? 0&1 0&1 0&1 0&1 0&1 0&1 1 0 0 ? ? 0 0 0 1 1 1 2 ? 1 0 0 ? ? ? 1 ? ? ? ? ? ? ? ? ? ? ? ? ? ? ? ? ? ? ? ? ? ? ? 0 ? ? ? ? 2 1 ? 0 1 1 1 1 0 ? ? ? ? ? ? ? ? ? ? ? ? ? ? ? ? ? ? ? ? ? ? ? ?

##### Apomorphy list

**Spinosauridae (with *Camarillasaurus*)**

Ch. 48: 0 → 1 Maxillary lateral teeth with circular midcrown cross section.

Ch. 92: 0 → 1 Middle and posterior caudal vertebrae with short prezygapophyses.

Ch. 119: 0 → 1 Tibia with a proximal diaphysis with length+width ratio greater than 2.

**Spinosauridae (minus *Camarillasaurus*)**

Ch. 49: 0 → 1 Maxillary lateral teeth with fluting (apico-basal ridges).

Ch. 94: 0 → 1 Anterior chevrons with longitudinal groove widened as a fossa.

**Baryonychinae**

Ch. 3: 0 → 1 V-shaped premaxilla-nasal suture

Ch. 30: 0 → 1 Square-shaped quadrate head

Ch. 57: 0 → 1 30 dentary teeth.

Ch. 64: 1 → 0 Middle cervical vertebrae with neural spines longer than tall

Ch. 74: 0 → 1 Dorsal vertebrae, posterior neural spines with basal webbing

Ch. 78: 0 → 1 Dorsal vertebrae with accessory centrodiapophyseal lamina

Ch. 97: 0 → 1 Coracoid with crescentic posterior process.

**Ceratosuchopsini (*Suchomimus* + *Ceratosuchops*)**

Ch. 26: 0 → 1 Frontal, postorbital facet depth more than 2/5 facet length.

Ch. 27: 0 → 1 Frontal with abrupt transition between the anterior half and the postorbital process

Ch. 40: 0 → 1 Basioccipital contribution to foramen magnum reduced, with exoccipitals closely placed.

**Spinosaurinae**

Ch. 63: 0 → 1 Middle and posterior cervical vertebrae with centra wider than tall

Ch. 66: 0 → 1 Anterior dorsal vertebrae with strongly developed hypapophysis

Ch. 69: 0 → 1 Anterior dorsal vertebrae with pneumatic foramen smaller than parapophysis

Ch. 71: 0 → 1 Middle and posterior dorsal vertebrae with deeply excavated prezygopara-diapophyseal fossa (prpadf).

Ch. 76: 0 → 1 Posterior dorsal vertebrae with neural spine base shorter than centrum

Ch. 77: 0 → 1 Middle and posterior dorsal vertebrae with reduced, knob-like parapophyses

Ch. 84: 0 → 1 Anterior caudal vertebrae with centro-costal laminae

Ch. 85: 0 → 1 Anterior caudal vertebrae with prezygo-costal fossa delimited by two laminae.

***Vallibonavenatrix* + (*Spinosaurus* + *Irritator*)**

Ch. 67: 0 → 1 Mid-dorsal vertebrae centra longer than 1.4 times centrum height

Ch. 72: 0 → 1 Middle and posterior dorsal centra with large pneumatic foramina

Ch. 83: 0 → 1 Anterior caudal vertebrae with well-marked spino-diapophyseal lamina

Ch. 91: 0 → 1 Shorter middle caudal centra.

***Spinosaurus* + *Irritator***

Ch. 109: 0 → 1 Ilium, orientation of pubic peduncle mostly anterior or ’kinked’ double facet with anterior and ventral components.

---

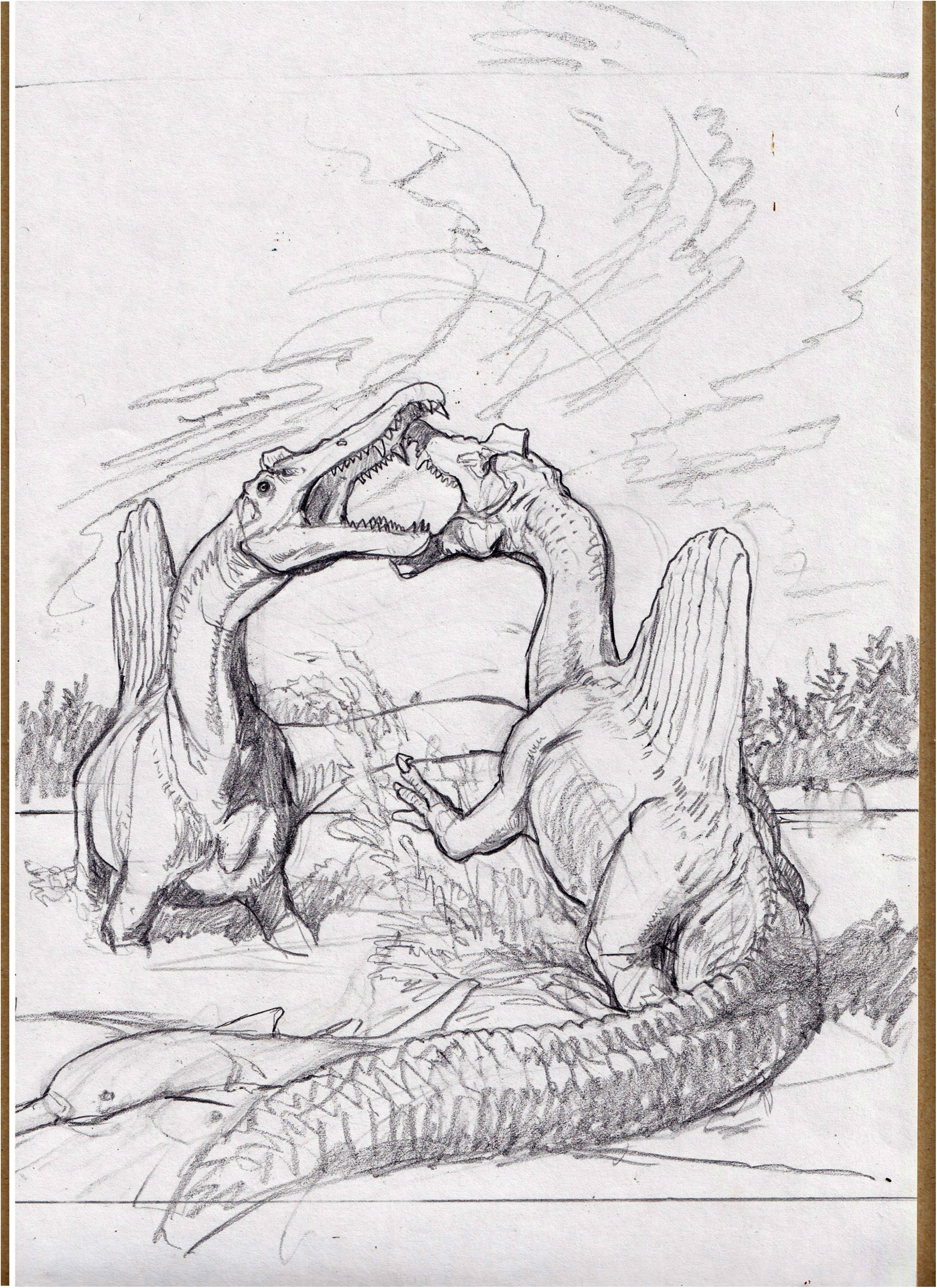

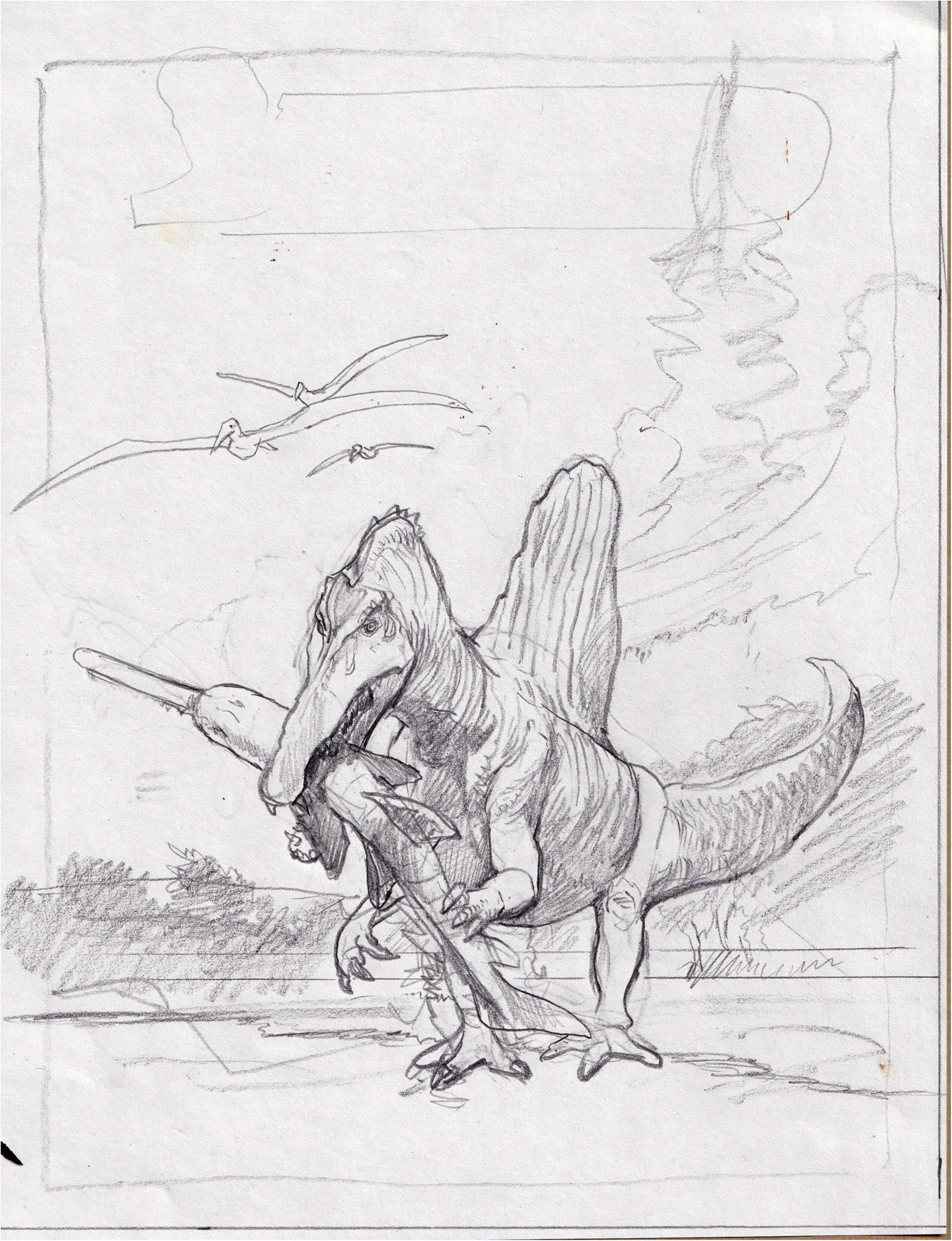

